# Bacterial meningitis in the early postnatal mouse studied at single-cell resolution

**DOI:** 10.1101/2023.01.11.523597

**Authors:** Jie Wang, Amir Rattner, Jeremy Nathans

**Author notes:** Address for editorial correspondence: Dr. Jeremy Nathans, 805 PCTB, 725 North Wolfe Street, Johns Hopkins University School of Medicine, Baltimore, MD 21205.

## Abstract

Bacterial meningitis is a major cause of morbidity and mortality, especially among infants and the elderly. Here we study mice to assess the response of each of the major meningeal cell types to early postnatal *E. coli* infection using single nucleus RNA sequencing (snRNAseq), immunostaining, and genetic and pharamacologic perturbations of immune cells and immune signaling. Flat mounts of the dissected arachnoid and dura were used to facilitiate high-quality confocal imaging and quantification of cell abundances and morphologies. Upon infection, the major meningeal cell types – including endothelial cells (ECs), macrophages, and fibroblasts – exhibit distinctive changes in their transcriptomes. Additionally, ECs in the arachnoid redistribute CLDN5 and PECAM1, and arachnoid capillaries exhibit foci with reduced blood-brain barrier integrity. The vascular response to infection appears to be largely driven by TLR4 signaling, as determined by the nearly identical response induced by LPS administration and by the blunted response to infection in *Tlr4^-/-^* mice.

## Introduction

The brain and spinal cord are protected, both physically and immunologically, by the meninges, a multi-layered tissue that occupies the space between the CNS parenchyma and the surrounding bone (Figure 1A; Coles et al., 2017a; Weller et al., 2018). Starting from the surface of the brain and moving toward the skin, the meninges consists of: (1) the pia, a thin and semi-permeable layer of cells that allows passage of small molecules and proteins between the CNS parenchyma and the cerebrospinal fluid (CSF), (2) the sub-arachnoid space, a highly vascularized region containing fibroblasts and immune cells that is filled with CSF and supported by a web of trabeculae (the name “arachnoid” reflects its spiderweb-like structure), (3) the arachnoid mater, an epithelial layer that serves as the outer boundary of the CSF-accessible space, and (4) the dura, a fibrous layer containing draining sinuses (veins), lymphatics, fibroblasts, and immune cells.

**Figure 1.**
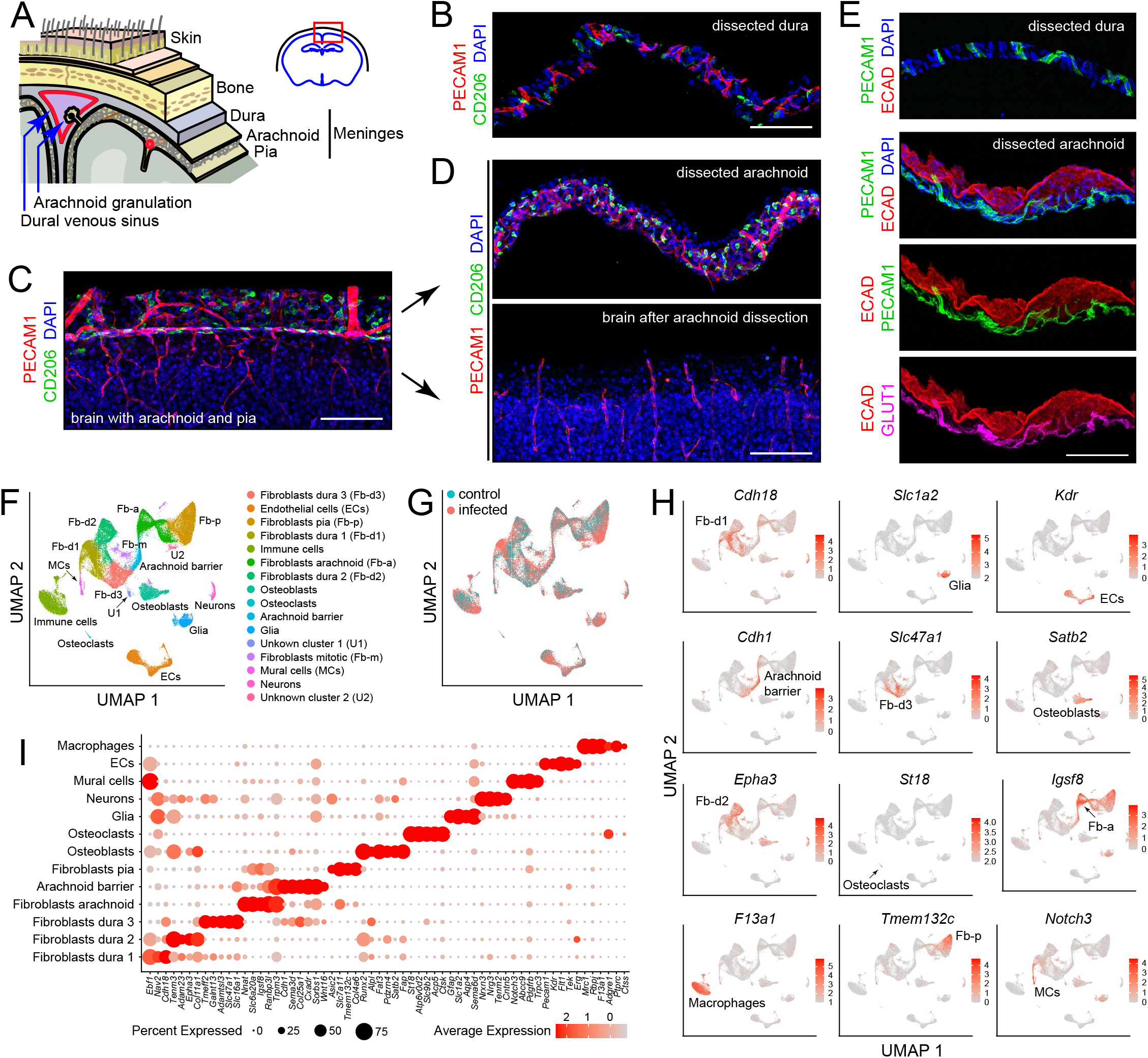
Dissection and single nucleus (sn) RNAseq of mouse meninges. (A) Diagram of the tissue layers between brain and skin, corresponding to the red rectangle in the coronal section through the brain and skull (upper right). (B) Cross-section of dissected dura stained for PECAM1 (endothelial cells) and CD206 (macrophages). (C) Coronal section through cortex (lower) and overlying pia and arachnoid (upper) stained for PECAM1 and CD206. (D) Cross-scetion of the isolated arachnoid (upper panel) and the denuded brain (lower) stained for PECAM1 and CD206. (E) Dissected dura and arachnoid stained for PECAM1 and ECAD (arachnoid epithelium); arachnoid is also stained for GLUT1 (a BBB marker; bottom panel). (F) UMAP plot of combined control and infected meninges snRNAseq datasets with cell clusters differentially colored and labeled. (G) UMAP plots of separated control and infected meninges snRNAseq datasets. (H) UMAP plots, as in panel (F) showing transcripts that are highly enriched in each of 12 cell clusters (labels in each panel). (I) Dot plot showing some of the transcript abundances that most clearly discriminate among major meningeal cell types, as well as contaminating neurons and glia. Scale bars: B-E, 100 um. All tissue and data in this and other figures are from P6 mice. The immunostaining and histochemical probes in this and other figures are indicated adjacent to the corresponding panel(s), with lettering color-coded to match the corresponding fluorescent color.

The meninges hosts a diverse collection of immune cells, including macrophages [referred to as barrier-associated macrophages (BAMs) or CNS-associated macrophages (CAMs)], monocytes, innate lymphoid cells (ILCs), T-cells, B-cells, dendritic cells, and mast cells. Recent single cell RNA sequencing (scRNAseq) and immuno-phenotyping have revealed distinctive layer-specific types and abundances of these immune cells (Rua and McGavern, 2018; Alves de Lima et al., 2020). Macrophages are especially abundant, and they can be divided into several molecularly distinct classes that are characterized by different half-lives and capacities for self-renewal (Mrdjen et al., 2018; Kierdorf et al., 2019; Van Hove et al., 2019; Masuda et al. 2022). Layer specific diversity is also observed among meningeal fibroblasts, with molecularly distinctive pial, arachnoid, and perivascular fibroblasts, as well as two types of dural fibroblasts (DeSisto et al., 2020; Derk et al., 2021).

A wide variety of CNS injuries and disease processes involve the meninges, including stroke, traumatic brain injury, neuroinflammatory conditions such as multiple sclerosis, neurodegenerative diseases, and infections (Derk et al., 2021; Alves de Lima et al., 2020). Disease and injury responses have been observed among meningeal immune cells, fibroblasts, and vasculature (Rua and McGavern, 2018; Derk et al., 2021). Recent experiments with animal models imply that some of these changes play a causal role in injury or disease pathology. For example, in mouse models of stroke, genetic ablation of meningeal mast cells reduces infarct size and brain swelling (Arac et al., 2014).

The oldest and best-established pathophysiologic role for the meninges is as a site of bacterial, fungal, or viral infection (Uiterwijk and Koehler, 2012; Williamson et al. 2017; Kohil et al., 2021). Bacterial meningitis is most common among young children and the elderly, and it is generally initiated by a blood-borne infection (Ku et al., 2015; McGill et al., 2016). The annual incidence of bacterial meningitis ranges from ~2 per 100,000 people in Western Europe and North America to 100-1,000 per 100,000 people in the Sahel region of Africa (Ku et al., 2015; GBD 2016 Meningitis Collaborators, 2018). In Western Europe and North America, mortality from bacterial meningitis is 10-20%, with >30% of survivors experiencing residual neurologic defects (McGill et al., 2016; Eisen et al., 2022). The molecules and mechanisms that mediate bacterial adhesion to and invasion of the meningeal vasculature are objects of active investigation (Coureuil et al., 2017).

Meningitis in the neonatal period, when the immune system is immature, typically results from bacterial infection during or immediately prior to delivery. Bacteria, most commonly Group B *Streptococci* and *E. coli*, are introduced into the bloodstream through breaks in the infant’s skin or via intra-amniotic infection (Gaschignard et al., 2011; Shane et al., 2017). The incidence of bacterial meningitis in neonates is ~0.3 per 1,000 live births in developed countries and ~4 per 1,000 live births in less developed countries (Ku et al., 2015). Among survivors of neonatal meningitis, 20-70% (the number depending on the country) are left with long-lasting neurologic sequelae, including hearing loss, epilepsy, and learning and/or behavioral disabilities (Peltola et al. 2021).

In the present work, we describe the molecular and cellular responses of meningeal cells in a mouse model of neonatal *E. coli* meningitis. Responses to infection were observed in every major meningeal cell type, including endothelial cells (ECs), macrophages, and fibroblasts. We have also used genetic and pharmacologic perturbations of immune cells and pathways to explore communication networks responsible for this complex multi-cellular response.

## Results

### Single nucleus sequencing and flatmount imaging of the mouse meninges

Our point of departure in studying the murine meninges was to utilize a simple protocol for dissecting the arachnoid and the dura free from adjacent tissues. When the skull and brain are separated in the absence of fixation, the natural cleavage plane is between the arachnoid and dura (Figure 1A). [In the text that follows, we will refer to the arachnoid mater and the sub-arachnoid space collectively as “the arachnoid”.] The arachnoid can then be peeled from the brain surface and the dura can be peeled from the inner surface of the skull. Immunostaining of the isolated arachnoid, the isolated dura, and the denuded brain reveals vasculature (expressing PECAM-1/CD31) and macrophages (expressing CD206/MRC1) in both arachnoid and dura (Figure 1B-D). The arachnoid epithelium expresses E-cadherin (ECAD). Arachnoid ECs, but not dura ECs, express GLUT1, a blood-brain barrier (BBB) marker (Figure 1E). These and all other analyses in this study were conducted at postnatal day (P)6.

For transcriptome analyses at cellular resolution, we sought to obtain as representative a sampling of cell types as possible and to minimize RNA synthesis or degradation after dissecting the dura and arachnoid. Therefore, we avoided enzymatic tissue dissociation and, instead, purified nuclei following tissue homogenization. The meninges were obtained from P6 mice, either without infection (“control”) or following a single subcutaneous injection of *E. coli* K1 at P5 (“infected”). (All experiments, except for those shown in Figures 6C-F, 7, and 8E were conducted with FVB/NJ mice.) Single nucleus (sn)RNAseq data were obtained from 14,356 and 34,585 nuclei from control and infected mice, respectively, with a mean of 1,320 transcripts sequenced per nucleus, using the 10X Genomics Chromium platform (Supplemental Table 1). Pairwise Pearson correlations among the five snRNAseq datasets (two control and three infected) shows correlations of 0.98-1.00 within the same group and 0.90-0.93 between infected vs. control groups (Supplemental Figure 1).

Fourteen principal cell clusters were identified with Seurat, and their identities were assigned by immunostaining and with reference to published data, as seen in the Uniform Manifold Approximation and Projection (UMAP) plots in Figure 1F and 1H (Supplemental Table 2). One cluster, labeled “immune cells”, consists largely of macrophages but also encompasses other immune cells, as described in detail below. Two clusters represent neurons and glia, presumably brain contaminants. One small cluster derives from mitotic fibroblasts, and two small clusters (U1 and U2) are unidentified. Strikingly, five large clusters represent fibroblasts – one pial, one arachnoid, and three dural. A dot plot of 61 transcripts that exhibit cell-type-specific patterns of enrichment supports these cell cluster assignments and also illustrates a pattern of partial overlap in gene expression among the five fibroblast clusters (Figure 1I). Comparing UMAP plots of control and infected datasets reveals shifts in the positions of the immune cell and fibroblast clusters with infection (Figure 1G).

Flatmounts of the isolated arachnoid and dura permit confocal imaging across the full depth of each of these tissues (Supplemental Figure 2). Additionally, the dura can be imaged as a flatmount while it remains attached to the inner surface of the skull, although this arrangement reduces tissue access to antibody and washing solutions. Arachnoid flatmounts show a high density of macrophages marked by (1) co-expression of macrophage markers CD206 and LYVE1, which localize to distinct cytoplasmic/surface compartments, and (2) expression of PU.1, which localizes to the nucleus (Supplemental Figure 2A-C). These cells, and other nonmacrophage immune cells, also express CD45/PTPRC, a general marker for hematopoietic cells (Supplemental Figure 2A-C). Flatmounts of the peripheral dura (i.e., outside the sinuses) show elongated perivascular macrophages expressing CD206 and LYVE1, and additional immune cells expressing CD45 (Supplemental Figure 2D).

The vasculature can be visualized in arachnoid and dura flatmounts by immunostaining for PECAM-1, and, in arachnoid flatmounts, by immunostaining for tight junction markers Occludin (OCLN), Zonula occludens-1 (ZO-1), and Claudin-5 (CLDN5) (Supplemental Figure 2C and below). In dura flatmounts with the bone attached, perivascular fibroblasts express FOXP2, and osteoblasts express SATB2 (Supplemental Figure 2D).

### Responses of meningeal immune cells to infection

In comparing control vs. infected data sets for each of the principal cell clusters, scatter plots encompassing all transcripts reveal relatively few changes in the contaminating neuron and glia clusters and many more changes in each of the principal meningeal clusters, with roughly equal numbers of up- and down-regulated transcripts (Supplemental Figure 3 and Supplemental Table 3). Dot-plot comparisons show several transcripts – for example, *Kiz, Slc39a14, Ap4e1*, and *Cp* – that are up-regulated across nearly all clusters (Supplemental Figure 4A). Changes in other transcripts were limited to smaller subsets of cell clusters, for example, the down-regulation of *Col14a1, Col8a1*, and *Slc4a10* in dural fibroblasts, the down-regulation of *Igsf8, Tmtc4*, and *Zfp536* in arachnoid and pial fibroblasts and arachnoid barrier cells, and the up-regulation of *Cxcl2* in macrophages (Supplemental Figure 4A and B). Hierarchical clustering in a gene set enrichment analysis (GSEA) with control vs. infected samples for the principal meningeal cell types shows prominent induction of inflammatory responses (“complement”, “interferon gamma response”, “IL6 JAK STAT3 signaling”, “TNFA signaling via NFKB”) across all cell types and a reduction in apical barrier (i.e. tight junction) transcripts in ECs and arachnoid barrier cells (Supplemental Figure 5).

The *E. coli* cells used for infection express Red Fluorescent Protein (RFP), which facilitates their visualization in tissue. At P6, one day after subcutaneous inoculation, *E. coli* cells were typically observed in a patchy distribution in the arachnoid and dura, and at much sparser density within the brain (Figure 2A-D). The spatial heterogeneity in *E. coli* accumulation within the meninges, together with animal-to-animal variation in the severity of infection, likely accounts for some degree of variability in the cellular alterations associated with infection. To address this issue, representative images were chosen for the figures and for quantification.

**Figure 2.**
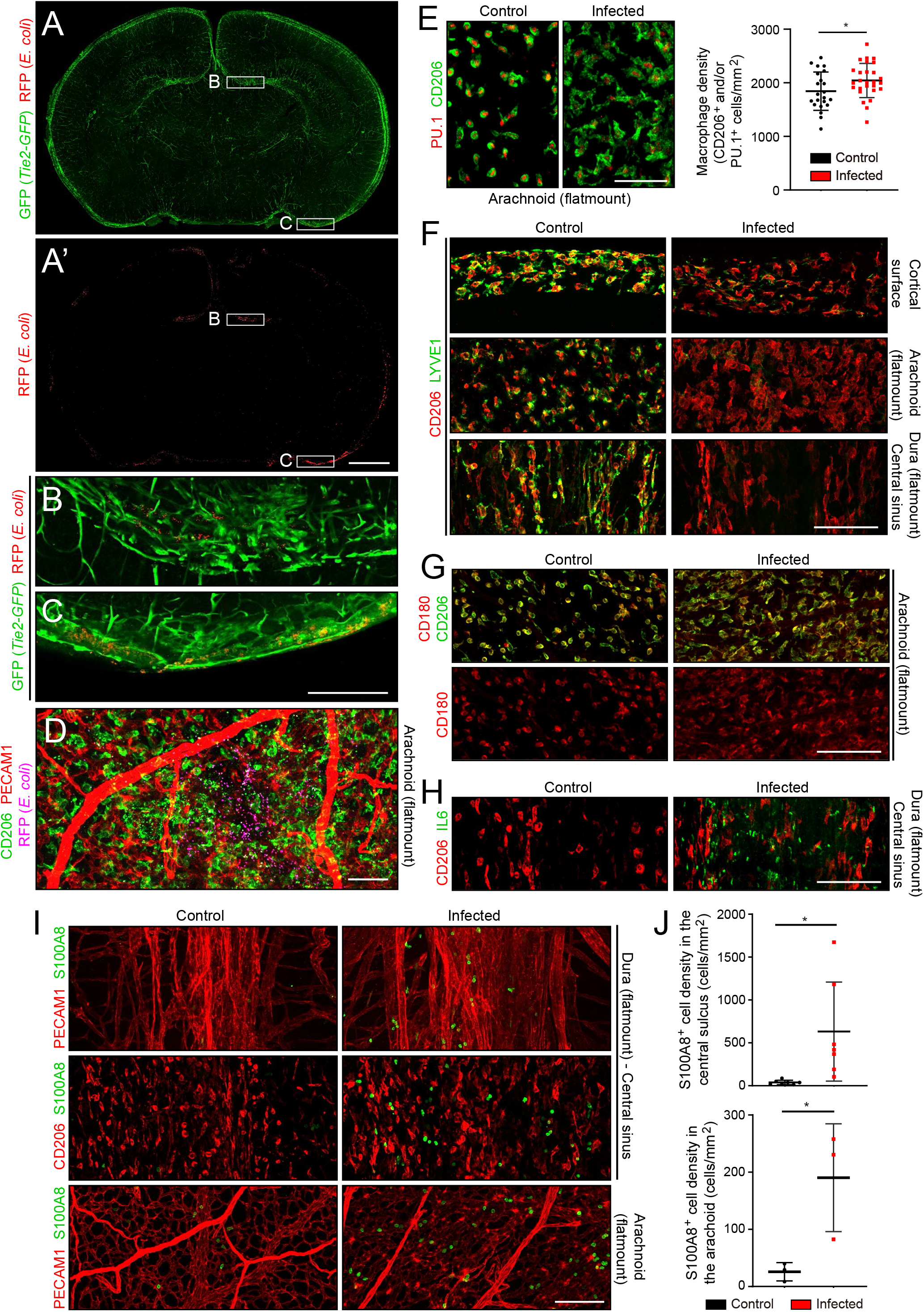
The *E. coli* meningitis model and some immune cell responses. (A-C) Coronal sections of P6 brain with arachnoid, one day after a subcutaneous injection of 1.2 x10^5^ RFP-expressing *E. coli* K1. Regions within the rectangles labeled (B) and (C) are enlarged below. The *Tie2-GFP* transgene is expressed in ECs. (D) Arachnoid flatmount showing scattered *E. coli* (RFP; false colored magenta). (E) Arachnoid macrophages, visualized with nuclear immunostaining for transcription factor PU.1 and cytoplasmic staining for CD206, show cytoplasmic enlargement upon infection (left), but only a small increase in number (right). (F) Infection leads to a selective reduction in LYVE1 immunostaining, and little or no change in CD206 immunostaining, in macrophages in the arachnoid and dura. (G) Infection leads to little or no change in CD180 and CD206 immunostaining in macrophages in the arachnoid. (H) IL6 increases in dural fibroblasts in the central sinus. (I and J) Increase in cells immunostained for S100A8 in the arachnoid and dura of infected mice. All infected tissue and data in this and other figures are from P6 mice that had been infected 22 hours earlier. Scale bars: A and A’, 500 um; B-I, 100 um. In this and all other figures: n.s., not significant (i.e. p>0.05); *, p <0.05; **, p < 0.01; ***, p < 0.001; ****, p < 0.0001.

Despite the presence of *E. coli*, the density of macrophages in the arachnoid, which are normally present at ~2,000 cells per mm^2^, increased by only ~10% (Figure 2E). However, macrophage morphology changed dramatically with infection, from small and rounded to large and irregularly shaped (Figure 2E). Infection also led to a reduction in LYVE1 abundance, but no change in CD180 or CD206 abundance, in macrophages (Figure 2F and G). In the dura of infected mice, IL6 was induced in fibroblasts (Figure 2H), and, in both the dura and arachnoid, the number of cells expressing S100A8, a marker for monocytes and immature macrophages, increased ~10-fold (Figure 2I and J).

For a more comprehensive assessment of the immune response to infection, we further parsed the immune and immune-related clusters into microglia (a brain contaminant), innate lymphoid cells/T cells (ILC/T), osteoclasts (a contaminant from the skull), monocytes (MCs) and monocyte-derived cells, resident macrophages (MPs; subdivided by CCL2 expression), and inflammatory macrophages [subdivided by IL1 receptor type 1 (IL1R1) expression] (Figure 3A-C, and 3D, lower panel; Supplemental Table 2). Although the relative abundances of the principal meningeal cell types did not change with infection, as judged by counting nuclei in the five snRNAseq libraries (Figure 3D, upper panel), parsing the individual immune cell types revealed a ~2-fold decrease in the abundance of CCL2^-^ resident macrophages, a >10-fold increase in inflammatory macrophages (IL1R1^+^ and IL1R1^-^), and a several-fold increase in the abundance of monocytes or monocyte-derived cells (Figure 3D, lower panel). The changes in macrophage abundances could be largely explained if ~50% of the CCL2^-^ resident macrophages present prior to infection converted to inflammatory macrophages with infection. Among genes that are either up- or down-regulated in CCL2^-^ or CCL2^+^ resident macrophages, several dozen show greater than 5-fold changes in abundance (Figure 3E and Supplemental Table 4).

**Figure 3.**
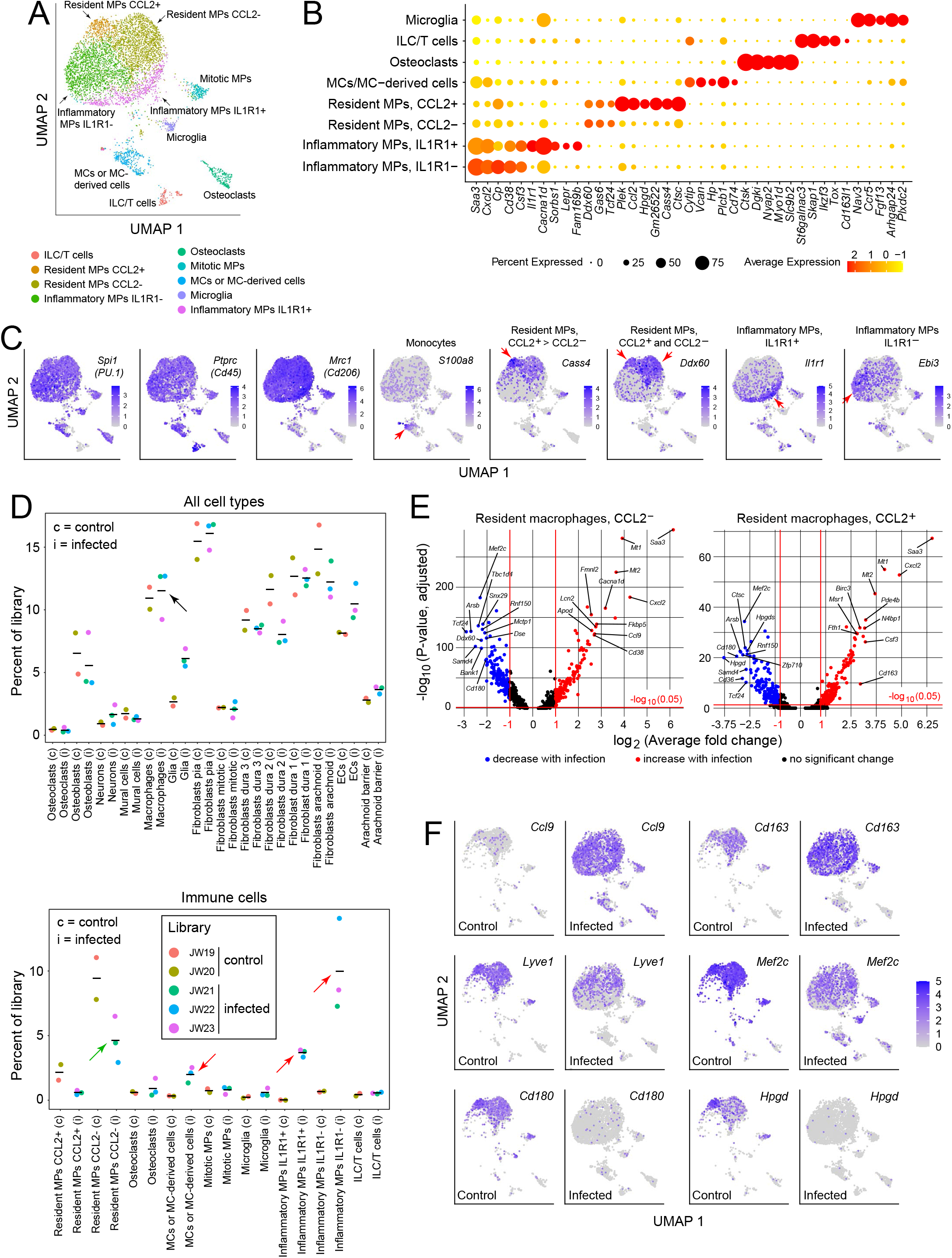
Immune subtypes and their responses to infection. (A) snRNAseq UMAP plot for immune cells from combined control and infected meninges. (B) Dot plot showing some of the transcript abundances that most clearly distinguish among meningeal immune cells. (C) UMAP plots, as in panel (A) showing eight transcripts that are expressed by all (left three panels) or by distinct subsets (right five panels) of macrophage subtypes and macrophage-like cells. Red arrows highlight regions within the UMAP clusters that correspond to distinct cell types, as defined in panel (A). (D) Comparing the number of nuclei in control (c) vs. infected (i) snRNAseq datasets across all meningeal cell types (upper panel) and across immune cells (bottom panel). The fraction of cells comprising the general category “macrophages” shows no change with infection (black arrow in upper panel). However, the lower panel shows that several macrophage subsets decrease (green arrow) or increase (red arrows) in abundance with infection. (E) Volcano plots for CCL2^-^ and CCL2^+^ macrophages showing control vs. infected snRNAseq transcript abundances (see Supplemental Table 4). (F) UMAP plots, as in (A), comparing control vs. infected snRNAseq for six genes.

The UMAP plots in Figure 3F compare the expression levels of six genes in control vs. infected immune cells and they illustrate the appearance of inflammatory macrophages, represented by the lower ~50% of the macrophage cluster, specifically in the infected meninges (see also Figure 3A). Figure 3F also illustrates the diversity of macrophage transcript changes with infection, with dramatic up-regulation of *Ccl9*, modest up-regulation of *Cd163*, modest down-regulation of *Lyve1* and *Mef2c*, and dramatic down-regulation of *Cd180* and *Hpgd*. Interestingly, by immunostaining, CD180 levels showed little change at this time point (Figure 2G), suggestive of a long protein half-life, whereas LYVE1 levels showed a large reduction (Figure 2F), suggestive of post-translational as well as transcriptional down-regulation.

### Responses of meningeal fibroblasts to infection

Fibroblasts constitute the most abundant cell type in the arachnoid and dura (Figure 1F, Supplemental Table 1, and Supplemental Figure 6A), and each of the five meningeal fibroblast subtypes shows numerous changes in transcript abundances in response to infection (Supplemental Figures 3–8 and Supplemental Table 3). Here, we highlight two gene families, collagens and SLC transporters, in which multiple family members show expression changes in control vs. infected meningeal fibroblasts. Transcripts coding for multiple collagen subtypes are down-regulated by infection: among the 50 members of the collagen gene family, 25 show detectable expression in the meninges by snRNAseq and 19/25 are down-regulated but only 2/25 are up-regulated (Supplemental Figure 7). Two examples are shown in the UMAP plots in Supplemental Figure 6B: *Col14a1* is down-regulated in type 1 and 2 dura fibroblasts, and *Col25a1* is down-regulated in arachnoid barrier cells and type 3 dura fibroblasts. Similarly, multiple transcripts coding for SLC transporters are down-regulated in meningeal fibroblasts. Of the ~350 members of the mouse *Slc* gene family with detectable expression in meningeal cells, 37 show infection-dependent changes in transcript abundance in one or more meningeal cell types by snRNAseq, with 21/37 down-regulated and 6/37 up-regulated by log_2_-fold >0.25 following infection (Supplemental Figure 8). The UMAP plots in Supplemental Figure 6B show down-regulation of *Slc16a1* in dura fibroblasts. *Slc39a14*, which codes for a divalent metal transporter, is unusual in its substantial up-regulation with infection (Supplemental Figure 8B).

In contrast to the pattern of down-regulation among the majority of *Col* and *Slc* transcripts, across the full transcriptome similar numbers of transcripts are up- and down-regulated by infection in each of the major meningeal cell types (Supplemental Figure 3). Among fibroblasts, examples of transcripts that are up-regulated include *Alk* in arachnoid fibroblasts, *Scara5* in pial fibroblasts, *Nrg3* in type 1 dura fibroblasts, *Camk4* in type 2 dura fibroblasts, and *Lbp* in type 3 dura fibroblasts (Supplemental Figure 6B). Additionally, infection up-regulates IL6 protein and *Il6* transcripts in fibroblasts (Figure 2H and Supplemental Figure 6B).

### Responses of meningeal vasculature to infection

In the context of bacterial meningitis, meningeal ECs serve as both a portal of entry for bacteria and immune cells and as a site of pathologically increased vascular permeability (Kim et al., 1997; Barichello et al., 2013; Coureuil et al., 2017). To explore the meningeal EC response to infection, we first parsed the EC UMAP into clusters derived from arachnoid, dura, and arterial ECs based on published markers (e.g. BBB vs. non-BBB markers for arachnoid vs. dura ECs, respectively; Figure 4A; Supplemental Tables 2 and 5). Infection leads to an expansion in the size of the arachnoid and dura EC clusters (Figure 4A and E), indicative of increased heterogeneity in transcriptome content, with little or no change in the proportions of the different EC subtypes (Figure 4B). Multiple transcriptome changes distinguish the infection responses of different EC clusters, including down-regulation of transcripts coding for amino acid transporters SLC7A1 and SLC7A5 in arachnoid ECs and down-regulation of transcripts coding for plasmalemma vesicle associated protein (PLVAP; a marker of high permeability vasculature) and the mechanosensory channel PIEZO2 in dura ECs (Figure 4C-E). Transcripts coding for VEGFR2/KDR are down-regulated in both arachnoid and dura ECs (Figure 4C-E).

**Figure 4.**
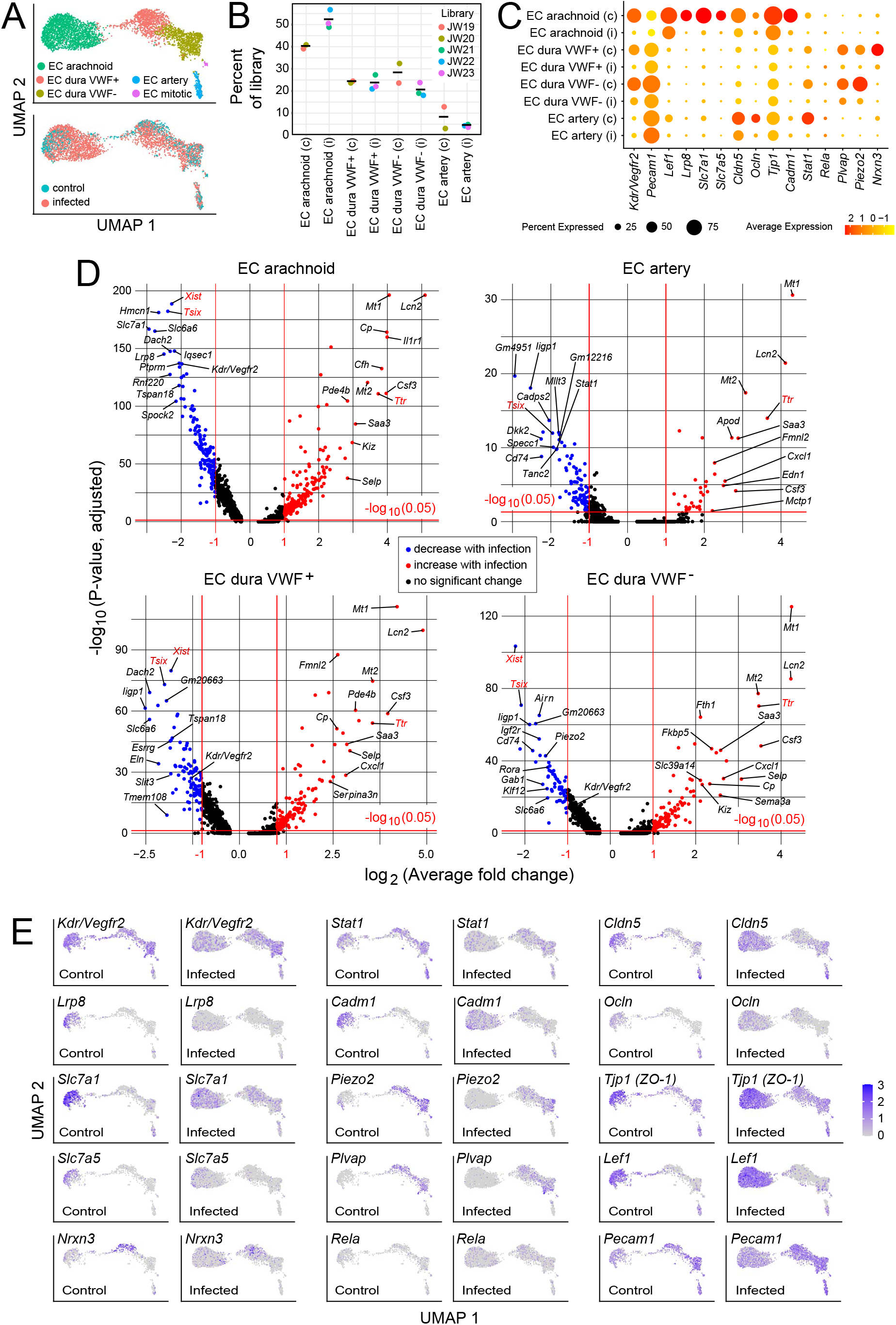
Changes in EC gene expression with infection. (A) snRNAseq UMAP plots for ECs from combined control and infected meninges. (B) The number of nuclei from different EC subtypes is consistent across the five snRNAseq libraries (control: JW19, JW20; infected: JW21-JW23). (C) Dot plot showing changes in transcripts abundances in EC subtypes in control (c) vs. infected (i) snRNAseq datasets. (D) Volcano plots for four EC subtypes showing control vs. infected snRNAseq transcript abundances. *Xist* and *Tsix* transcripts, referable to sex differences among embryos, are marked in red. *Ttr* transcripts, also marked in red, likely represent contamination from choroid plexus RNA and are present in 2/3 infected snRNAseq samples. (E) UMAP plots, as in (A), comparing control vs. infected snRNAseq for 15 genes.

In arachnoid flatmounts, infection produces mislocalization and clustering of CLDN5 and PECAM1, disorganized capillary morphology, and an expansion of the area covered by capillaries (Figure 5A-C, and Supplemental Figure 9). Immunoblotting of arachnoid proteins shows a modest reduction in the mean level of CLDN5, but this trend did not reach statistical significance due to the relatively high animal-to-animal variability in the infected group (Figure 5D). Functionally, there are patchy deficiencies in vascular barrier integrity following infection, with a spatial distribution that closely matches the clustering of CLDN5 and PECAM1, as revealed by extravasation of Sulfo-NHS-biotin, a low molecular weight intravascular tracer (Figure 5A and B). The same infection-associated vascular phenotypes were also seen one day after an intraperitoneal (IP) injection of 10 mg/kg lipopolysaccharide (LPS), a potent activator of the innate immune response to gram-negative bacteria such as *E. coli* (Figure 5A-C).

**Figure 5.**
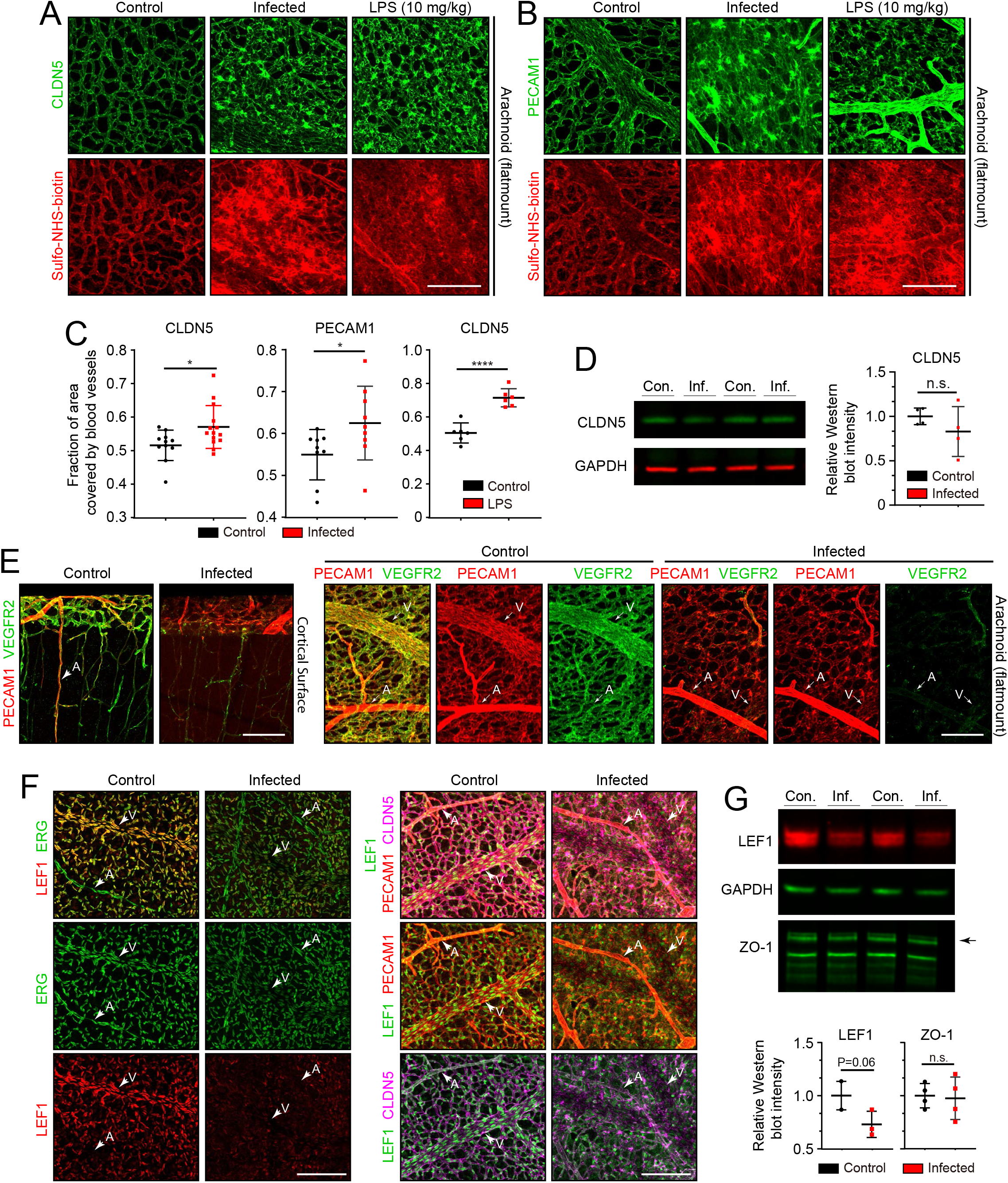
Changes in EC morphology and EC protein abundance and localization with infection. (A) CLDN5 localization and Sulfo-NHS-biotin leakage in arachnoid vasculature following infection or LPS administration. (B) PECAM1 localization and Sulfo-NHS-biotin leakage in arachnoid vasculature following infection or LPS administration. (C) Infection or 10 mg/kg LPS treatment increases the area covered by vasculature in flatmounts of arachnoid. (D) Immunoblotting shows a modest, but not statistically significant, reduction in CLDN5 level relative to GAPDH level in the arachnoid with infection. (E) Reduced KDR (VEGFR2) immunstaining in arachnoid ECs with infection. (F) Reduced EC nuclear LEF1 immunostaining in capillaries and veins, and reduced PECAM1 and CLDN5 staining in veins in the arachnoid with infection. (G) Immunoblotting shows a modest reduction in LEF1 level and no change in ZO1 level relative to GAPDH level in the arachnoid with infection. Scale bars: A and B, 100 um; E and F, 100 um.

Two well-studied signaling pathways are known to control CNS vascular permeability: VEGF and WNT. Consistent with the down-regulation of *Vegfr2/Kdr* transcripts seen in infected ECs by snRNAseq (Figure 4C-E), VEGFR2 immunostaining was also greatly reduced (Figure 5E). As VEGFR2 is the principal receptor for VEGF signaling in ECs, its down-regulation suggests that the enhanced vascular permeability associated with bacterial infection is not caused by increased VEGF signaling, a known mechanism for increasing vascular permeability (Senger et al., 1983; Roberts and Palade, 1995). WNT signaling in CNS vasculature maintains the bloodbrain barrier (BBB) and it is both mediated by and up-regulates the transcription factor LEF1, which binds to target genes in combination with beta-catenin (Sabbagh et al., 2018). In control arachnoid flatmounts, LEF1 accumulates in vein and capillary EC nuclei with minimal accumulation in arterial ECs (Figure 5F, left panels). Strikingly, in infected arachnoid flatmounts, LEF1 immunostaining in ECs is greatly reduced, but its expression persists in some non-ECs (Figure 5F, right panels). By immunoblotting, total arachnoid LEF1 levels are reduced by ~25% (Figure 5G). The right panels of Figure 5F also shows reductions in CLDN5 and PECAM1 specifically in arachnoid veins, which appear as “shadows” in the flatmount images. These data are consistent with a model in which infection leads to reduced BBB integrity in the arachnoid vasculature, at least in part, by reducing WNT signaling in ECs.

To assess the effects of infection on the organization of dura ECs, we took advantage of the large and unusually straight veins that occupy the dural sinuses adjacent to the skull’s sutures. Venous ECs throughout the body typically exhibit elongated nuclei that are aligned with the long axis of the vein, and therefore also with the direction of blood flow. With infection, this alignment is diminished in dural veins (Figure 6A and B), suggesting a general effect of infection on cytoskeletal organization within venous ECs. No assessment of permeability was made for the dura vasculature because the dura resides outside of the BBB territory (delimited by the arachnoid epithelial barrier) and, therefore, it exhibits high permeability in the control state.

**Figure 6.**
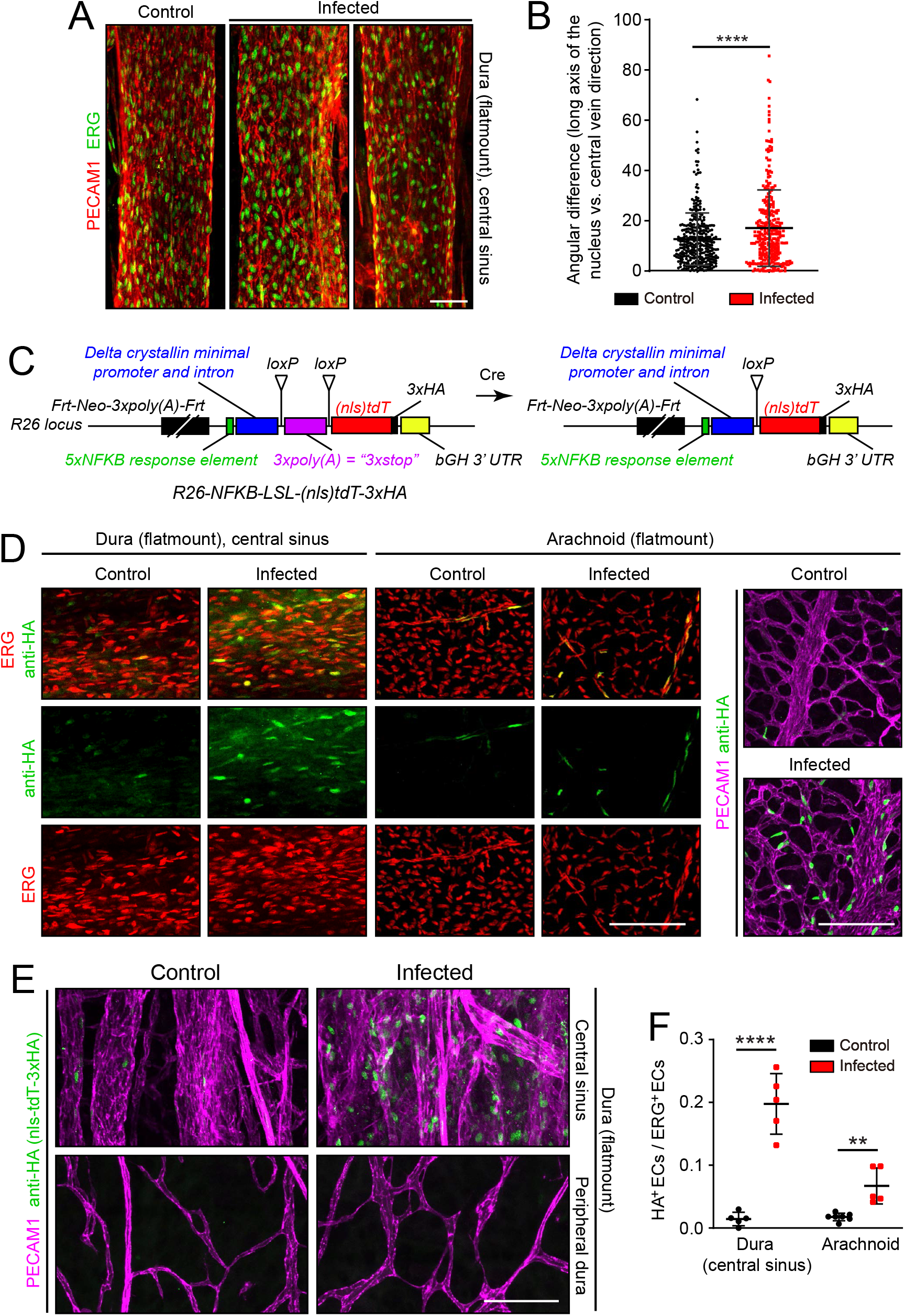
Infection causes disorganization of EC nuclear orientation in the dural venous sinus, and an increase in NF-kappa B signaling in ECs in the arachnoid and dura. (A) EC nuclei, visualized with ERG immunostaining, in the large vein of the central sinus. (B) Quantifying the orientation of the long axis of EC nuclei in the large vein of the central sinus, as shown in (A). (C) Structure of the NF-kappa B reporter before (left) and after (right) Cre-mediated recombination that removes a *loxP-transcription stop-loxP* (*LSL*) cassette. NF-kappa B reporter activation leads to expression of *nls-tdT-3xHA*. (D) Infection increases expression of the NF-kappa B reporter in a subset of ECs in the dura and arachnoid, as determined by immunostaining for HA. (E) In the dura, NF-kappa B reporter activation is observed in both ECs and non-ECs in the central sinus, but the NF-kappa B reporter is not activated in the peripheral dura. (F) Quantification of NF-kappa B reporter activation in ECs in the central sinus of the dura and in the arachnoid. Scale bars: A, 50 um; D and E, 100 um.

Microbial products, such as LPS, and any of a wide variety of cytokines could directly or indirectly alter EC structure and gene expression. As multiple immunologic stimuli are known to converge on the NF kappaB pathway and as the GSEA analysis implied that this pathway was induced upon infection (Supplemental Figure 5), we assessed NF kappaB signaling by generating reporter mice in which five tandem repeats of a canonical NF kappaB response element were inserted upstream of a minimal promoter to drive expression of a nuclear localized and 3xHA epitope-tagged tandem dimer Tomato (*nls-tdT-3xHA*) (Figure 6C). The parental version of this mouse line has the additional feature that a *loxP-transcription stop-loxP* cassette separates the promoter and the *nls-tdT-3xHA* coding region, permitting cell-type specific readout of NF kappaB signaling when crossed to a cell-type specific *Cre* transgene or knock-in allele. For the present analyses, we have used germline Cre-recombination to generate an allele that is permissive for reporter expression in any cell type.

In both dura and arachnoid, NF kappaB reporter expression was induced by infection almost exclusively in ECs, with reporter+ EC nuclei increasing from ~1% to ~20% in the dura and from ~1% to ~5% in the arachnoid. EC nuclei were identified based on ERG immunostaining (Figure 6D-F). A surprising feature of NF kappaB reporter expression in ECs is its spatial heterogeneity, with reporter expressing ECs adjacent to non-expressing ECs in an apparently random pattern. These data suggest cell-to-cell heterogeneity in meningeal EC responses to infection and are consistent with the observed broadening of infected EC clusters in the UMAP plots in Figure 4A and E.

### Genetic and pharmacologic perturbations of the immune response

To explore the role of specific immune pathways in the vascular changes associated with infection, we applied the *E. coli* infection paradigm to mice with null mutations in (1) *Tlr4*, the gene coding for the innate immune system’s LPS receptor, or (2) *Ccr2*, the gene coding for one of the receptors for monocyte chemoattractant protein-1 (MCP1/CCL2), a chemokine that recruits immune cells to sites of infection. In response to infection, the arachnoid vasculature of *Tlr4^-/-^* mice showed minimal changes in the distribution of CLDN5 and very few regions of Sulfo-NHS-biotin leakage. In contrast, *Ccr2^-/-^* mice showed a clumpy redistribution of CLDN5 and localized regions of Sulfo-NHS-biotin leakage (Figure 7A). Quantifying the area occupied by arachnoid vasculature showed that, in *Tlr4*^-/-^ mice, infection had essentially no effect, but, in *Ccr2^-/-^* mice, infection produced an increase similar to that produced in WT mice (Figure 7B). (For this experiment, the WT comparator strain is C57BL/6J, which matches the background of the *Tlr4^-/-^* and *Ccr2^-/-^* mice. The EC response to infection is milder in C57BL/6J mice compared to FVB/NJ mice.) In control (i.e., uninfected) *Tlr4^-/-^* and *Ccr2^-/-^* mice, the density of arachnoid macrophages was, respectively, ~70% and ~80% of the WT value, and this density rose by ~20% in infected *Tlr4*^-/-^ mice but showed little or no change in infected *Ccr2*^-/-^ or WT mice (Figure 7C).

**Figure 7.**
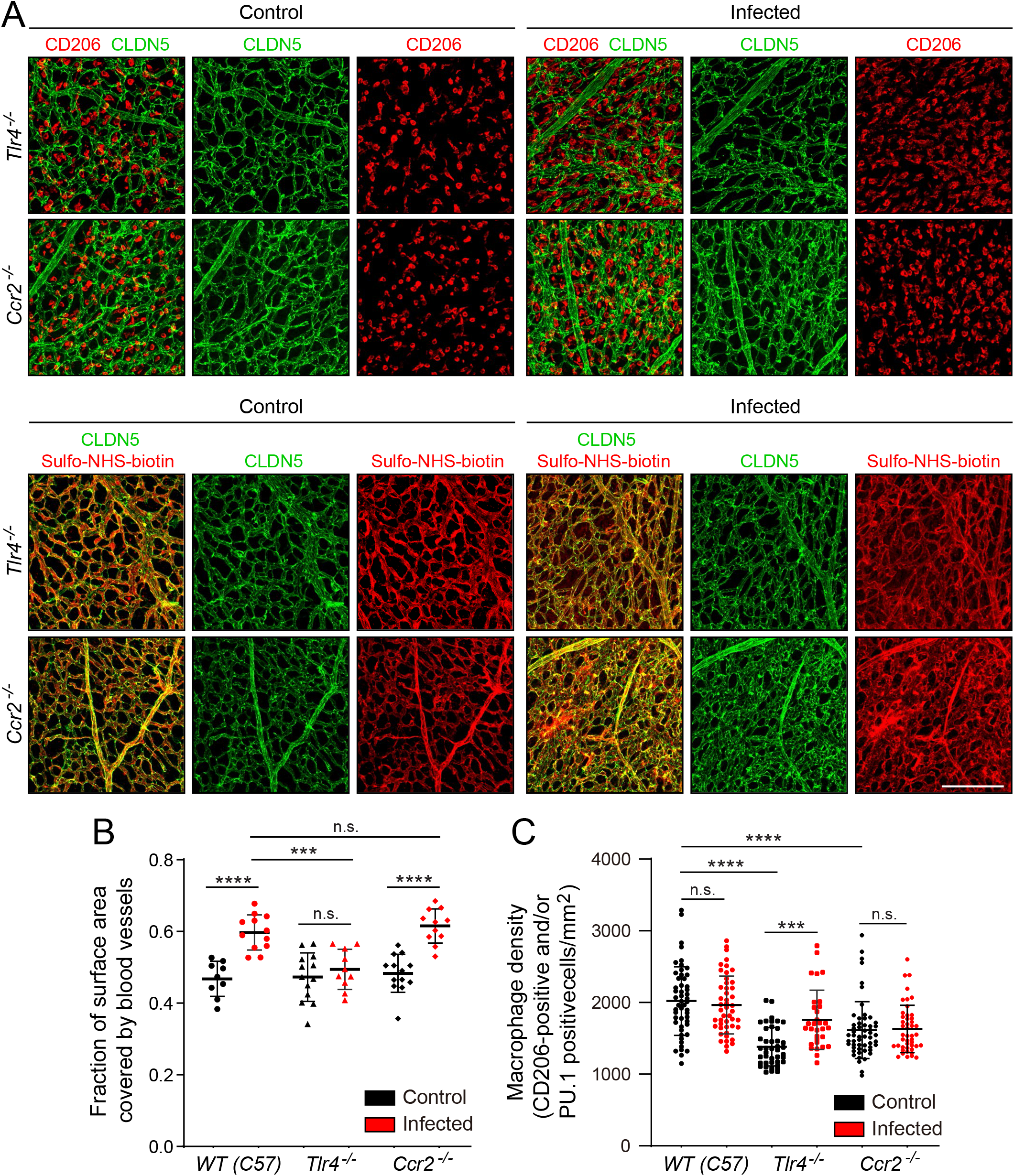
Effects of *Tlr4* KO and *Ccr2* KO on arachnoid EC responses to infection. (A) Arachnoid flatmounts of control vs. infected mice showing, in the upper panels, macrophage density (CD206) and vascular architecture (CLDN5) and, in the lower panels, vascular leakage (sulfo-NHS biotin). (B) Quantification of macrophage density in WT, *Tlr4^-/-^*, and *Ccr2^-/-^* arachnoid flat mounts in control vs. infected mice. (C) Quantification of vascular architecture in WT, *Tlr4^-/-^*, and *Ccr2^-/-^* arachnoid flat mounts in control vs. infected mice. Scale bar: A, 100 um.

As a complementary approach to gene inactivation, we used a single intra-cerebroventricular (ICV) injection of clodronate-containing liposomes to acutely and selectively eliminate arachnoid macrophages two days before *E. coli* inoculation (Figure 8A and C). Surprisingly, eliminating arachnoid macrophages had little or no effect on the infection-dependent redistribution of CLDN5 in arachnoid ECs, leakage of Sulfo-NHS-biotin, the increase in the area occupied by arachnoid vasculature, or the fractional increase in the number of ECs showing induction of the NF kappaB reporter (Figure 8B, D, and E). The principal differences between responses of mice receiving control liposomes vs. clodronate liposomes were the higher overall levels (i.e. both baseline and infected) of (1) the arachnoid blood vessel area (Figure 8D) and (2) the number of arachnoid ECs with NF kappaB reporter activation following clodronate treatment (Figure 8E).

**Figure 8.**
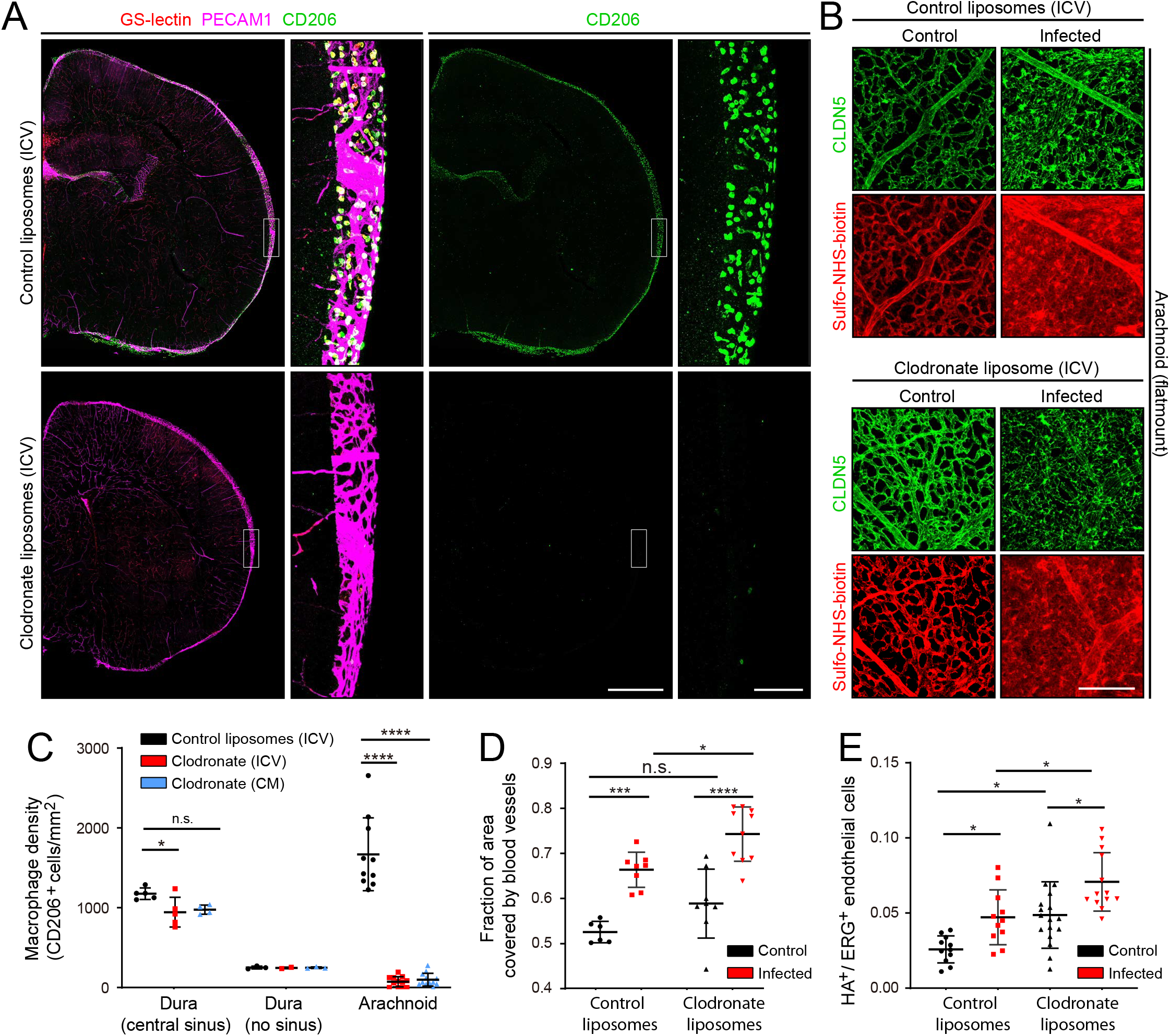
EC response to eliminating arachnoid macrophages with liposomal clodronate. (A) Intracerebroventricular (ICV) injection at P3 of empty liposomes vs. clodronate liposomes shows that clodronate almost completely eliminates arachnoid macrophages at P6, as visualized with CD206 immunostaining. (B) Little or no effect of macrophage elimination on CLDN5 relocalization and Sulfo-NHS-biotin leakage in arachnoid ECs in response to infection. (C) Quantification of arachnoid vs. dura macrophage abundance following ICV or cisterna magna (CM) injection of clodronate liposomes. (D) Quantification of vascular density in arachnoid flat mounts following injection at P3 with empty liposomes vs. clodronate liposomes in control vs. infected mice. (E) Quantification of NF-kappa B reporter activation in ECs in arachnoid flat mounts following injection at P3 with empty liposomes vs. clodronate liposomes in control vs. infected mice (Figure 6C-F). Scale bars: A, 500 um (low magnification) and 100 um (inset); B, 100 um.

Taken together, these experiments, together with the LPS treatment experiment (Figure 5A-C), show that (1) LPS stimulation of TLR4 signaling plays a central role in the response of the arachnoid vasculature to infection (CLDN5 and PECAM1 redistribution, vessel swelling, and leakage), and (2) this vascular response is largely independent of arachnoid macrophages, by far the most abundant immune cells in the arachnoid.

## Discussion

The present study defines the responses of cells in the mouse arachnoid and dura to bacterial meningitis in the early postnatal period. At this age, the immaturity of the adaptive immune system and the rapidity of infection imply that the host response depends largely, and perhaps exclusively, on the innate immune system. In response to bacterial infection, all of the major meningeal cell types – including ECs, macrophages, and fibroblasts – exhibit large and distinctive changes in their transcriptomes. In addition, ECs in arachnoid capillaries redistribute CLDN5 and PECAM1, arachnoid capillaries become enlarged and disorganized and they exhibit foci of reduced BBB integrity, and ECs in arachnoid capillaries and veins lose nuclear LEF1. These capillary responses to infection appear to be largely driven by TLR4 signaling, as determined by the response to LPS administration and by the blunting of these responses to bacterial infection in the absence of TLR4.

The role of LPS in mediating BBB breakdown has been intensively studied, primarily in the context of the vasculature within the brain parenchyma (Wispelway et al., 1988; Banks et al. 2015). In the brain, LPS treatment leads to a reduction in EC tight junctions secondary to a decrease in tight junction protein abundance and changes in tight junction protein localization (Peng et al., 2021). In multiple cell types, TLR4 signaling (i.e. LPS-induced signaling) activates the NF-kappa B pathway, and that connection presumably accounts for the NF-kappa B reporter activation in meningeal ECs observed here.

The NF-kappa B and WNT pathways can exhibit either positive and negative cross-regulation, depending on cell type and developmental context (Ma and Hottiger, 2016). By comparing BETA-CATENIN level, localization, and signaling in WT mouse embryo fibroblasts (MEFs) and in MEFs homozygous for inactivating mutations in IKKalpha or IKKbeta (the inflammation-activated kinases that phosphorylate the inhibitory binding partner of NF kappaB, leading to NF kappaB activation), Lamberti et al (2001) found that IKKalpha and IKKbeta phosphorylate BETA-CATENIN on different sites and with opposite effects. IKKalpha phosphorylation leads to BETA-CATENIN stabilization and increased WNT signaling, whereas IKKbeta phosphorylation leads to BETA-CATENIN destabilization and decreased WNT signaling. These precedents in other cell types suggest that reduced LEF1 – and presumably reduced canonical WNT signaling – in ECs in the infected arachnoid could reflect BETA-CATENIN downregulation via the activated NF kappaB pathway. Reduced canonical WNT signaling in ECs would be predicted to reduce BBB integrity (Rattner at al., 2022).

The flat geometry of the meninges and its superficial location between the brain and the skull present unusually favorable opportunities for both *in vitro* and *in vivo* microscopy. For *in vitro* analyses, the relative thinness of the dissected mouse arachnoid and dura provide excellent access to antibodies and also facilitate high quality confocal imaging without the need for chemical clearing agents, as described here. *In vivo*, thinned skull preparations that permit two-photon imaging of the mouse meninges have been described, and this approach can be applied to mice with fluorescent immune cells that have been inoculated with fluorescent bacteria to allow single-cell resolution in vivo imaging of bacterial meningitis in a native context (Kjos et al., 2015; Coles et al., 2017b; Manglani and McGavern, 2018). A largely unexplored opportunity also exists for *ex vivo* culture and live imaging of the dissected arachnoid and dura (Glimcher et al., 2008). Such preparations could permit high-resolution imaging of (1) bacterial movement across the vascular wall, (2) interactions between bacteria and immune cells, and (3) interactions between host cells. The use of fluorescent reporters of signaling pathway activity would further increase the value of such analyses (Kudo et al., 2018; Clark et al., 2021).

Despite decades of research, numerous gaps remain in our understanding of the pathophysiology of bacterial meningitis. These include: (1) the ways in which the imature immune system differs from the more mature immune system in its response to infection, (2) the relative importance and the precise roles of different immune cells and immune modulators, and (3) the roles played by changes in gene expression and cell behavior among non-immune cell types. In each of these areas, mouse models of meningitis, together with new technologies for interrogating these models, can provide insights that inform the understanding of human meningitis.

## Materials and Methods

### Mice

The following mouse lines were used: FVB/NJ (JAX#001800); C57BL/6J (JAX#000664); Tie2-GFP (JAX#003658); B6(Cg)-Tlr4tm1.2Karp/J (JAX#029015); B6.129S4-Ccr2tm1Ifc/J (JAX#004999); and *Rosa26-NF-kappaB* reporter mice (described below). All mice were housed and handled according to the approved Institutional Animal Care and Use Committee protocol of the Johns Hopkins Medical Institutions. Meninges snRNA-seq experiments and histological studies used postnatal day 6 (P6) mice with age-matched controls.

### Construction and genotyping of the NF-kappa B reporter

To construct the Cre-dependent reporter for NF-kappa B signaling at the *Rosa26* locus, the following elements were inserted (from 5’ to 3’ in the order listed) into a standard *Rosa26* targeting vector: an *frt*-phosphoglycerate kinase (*Pgk*)-neomycin (*Neo}-frt (FNF*) cassette, which includes a strong poly-adenylation signal; five tandem repeats of a canonical NF kappaB response element (GGGACTTTCC); a minimal Delta Crystallin promoter followed by an intron; a *loxP-transcription stop-loxP (LSL*) cassette; an open reading frame coding for a nuclear-localization signal-tdTomato-3xHA protein (*nls-tdT-3xHA);* and a bovine growth hormone 3’UTR. The *Rosa26-NF-kappaB-LSL-tdT* targeting construct with a 3’ flanking Diphteria toxin-A coding sequence was electroporated into R1 ES cells, which were then subjected to G418 selection. Clones harboring the targeted *Rosa26* locus were identified by Southern blot hybridization, karyotyped, and injected into blastocysts from Sv129 mice. Germline transmission to the progeny of founder males was determined by PCR. PCR primers for the parental allele: TGTCGGCCTGCAGCCAAAGCTTATCGA (sense, at the 3’ end of the *Neo* casette) and TGAAGTTCTCAGGATCGGTCGCTA (antisense, in the intron). PCR primers for the Cre-recombined allele: CCCCTCTGCTAACCATGTTCATGCCTT (sense, in the intron) and GGCAACCTTCCTCTTCTTCTTAGGCATGGTGG (antisense, at the 5’ end of the *nls-tdT-3xHA* open reading frame).

### Antibodies and other reagents

The following antibodies were used for tissue immunohistochemistry and immunoblotting: goat anti-CD45 (R&D Systems AF114-SP); rat anti-CD206/MRC1 (Bio-Rad MCA2235T); goat anti-CD206 (R&D Systems AP2535); rat anti-LYVE1 (Thermo Fisher/eBioscience 14-0443-82); rat anti-PU.1/Spi-1 (Novus Biologicals MAB7124); rat anti-CD180/RP105 antibody, PE (eBioscience 12-1801-81); goat anti-S100A8 (R&D Systems AF3059); rabbit anti-ERG (Cell Signaling Technologies 97249); rat anti-PECAM1/CD31 (BD Biosciences 553370); mouse anti-CLDN5, Alexa Fluor 488 conjugate (Invitrogen 352588); mouse anti-CLDN5 (Invitrogen 35-2500); rabbit anti-ZO1 (Invitrogen 40-2200); rabbit anti-LEF1 rabbit (Cell Signaling Technologies 2230); rabbit anti-LEF1, Alexa Fluor 647 conjugate (Cell Signaling Technologies 14022); rabbit anti-Occludin (Invitrogen 406100); goat anti-VEGFR2/KDR (R&D Systems AF644-SP); sheep anti-FOXP2 (R&D Systems AF5647-SP); rabbit anti-SATB2 (Abcam ab92446); rat anti-IL-6 (Biolegend 504501); rabbit anti-COL14A1 (Novus Biologicals NBP2-15940); chicken anti-GFP (Abcam Ab13970), rat anti-HA (Proteintech 7c9); rabbit anti-HA (homemade); goat anti-E-cadherin (R&D Systems AF748); rabbit anti-E-cadherin (Cell Signaling Technologies 3195); rabbit anti-AIFM3 (Novus Biologicals NBP1-76889); rabbit anti-COL25A1 (G-Biosciences ITT1021); goat anti-IGSF8 (R&D systems AF3117-SP); rabbit anti-NNAT (Abcam ab27266); mouse anti-GAPDH (Cell Signaling Technologies 97166S); rabbit anti-GAPDH (Cell Signaling Technologies 5174S); Streptavidin, Alexa Fluor 488 conjugate (Invitrogen S11223); Streptavidin, Alexa Fluor 647 conjugate (Invitrogen S32357). Alexa-Fluor-conjugated secondary antibodies were from Invitrogen. Infrared immunoblotting secondary antibodies were from LI-COR.

Other reagents used: Sulfo-NHS-biotin (Thermo Fisher Scientific #21217); LPS O111:B4 (Sigma-Aldrich L2630); Benzonase Nuclease, ultrapure (Sigma-Aldrich, E8263-5KU); clodronate and control liposomes: Mannosylated Macrophage Depletion kit (Encapsula NanoScience SKU#CLD-8914); Micro BCA Protein Assay kit (ThermoFisher Scientific 23235).

### *E.coli* infection and LPS injection

*E.coli* strain RFP-RS218 (O18:K1:H7) with K1 capsule is a clinical isolate from the CSF of a neonate with meningitis [Zhu et al., (2020); a generous gift from the late Dr. Kwang Sik Kim, Johns Hopkins Medical School, Baltimore, MD]. *E.coli* were grown overnight in Luria broth containing 100 μg/ml ampicillin at 37°C. The following day, 400 μL of the *E.coli* culture was added to 20 mL of fresh Luria broth for an additional 2 hours of culture at 37°C. The bacteria were washed one time in PBS, the OD at 620 nm was measured, and the concentration of the samples was adjusted prior to innoculation.

For most experiments, *E.coli* meningitis was induced in FVB mice. For experiments with *Tlr4^-/-^* and *Ccr2^-/-^* mice, C57BL/6J was used as the control to match the strain background of the KO lines. Briefly, a litter of P5 mice were randomly divided into two groups that were either subcutaneously injected in the back with 1.2 x 10^5^ CFU of *E.coli* RFP-RS218 in 20 μL PBS or not injected. Twenty-two hours later, the mice were sacrificed as described. *Tlr4*^-/-^ and *Ccr2*^-/-^ on a C57BL/6J background and *Rosa26-NF-kappaB* reporter mice on a mixed Sv129 x C57BL/6J background were subcutaneously injected with 8×10^4^ CFU of *E.coli* RFP-RS218. For LPS-injection, P5 mice were intraperitoneally injected with a single dose of LPS O111:B4 (10 mg/kg) or, for control mice, the same volume of PBS. Twenty-two hours later, the mice were sacrificed as described.

### Meningeal macrophage depletion with clodronate liposomes

CNS macrophages were depleted as described in Polfliet et al. (2001). Each mouse was injected with control or clodronate liposomes (3 μl of a 5 mg/ml stock solution) two days before being infected with *E. coli* or before receiving an LPS injection. Liposomes were allowed to warm from refrigerator temperature to room temperature for 1 hour prior to injection. P3 mice were deeply anaesthetized on ice and then slowly injected with liposomes using a Hamilton syringe (Hamilton Bonaduz, AG Switzerland) either in a lateral ventricle (intracerebroventricular administration) or in the cisterna magna.

### Tissue processing

To prepare isolated dura and arachnoid tissues for snRNAseq, immunoblots, and whole mount immunostaining, mice were deeply anesthetized on ice and then perfused via the cardiac route with PBS. The skullcaps (with dura mater attached) and brain (with arachnoid attached) were dissected in PBS. Using a fine tweezers, the arachnoid was gently peeled from the surface of the brain and either used for protein extraction, or immersion fixed in 2% paraformaldehyde (PFA)/PBS at room temperature for 1 hour for subsequent immunostaining. The dura was gently peeled from the skullcap and immersion fixed in 2% paraformaldehyde (PFA)/PBS at room temperature for 1 hour for subsequent immunostaining. Alternately, the skullcap and attached dura were immersion fixed overnight in 2% PFA/PBS at 4°C without further dissection. Following immersion fixation, each sample was washed three times in PBS. For vibratome sections, the brain and attached arachnoid was fixed overnight in 2% PFA/PBS at 4°C, washed the following day in PBS at 4°C for at least 3 hours, embedded in 3% agarose, and sectioned at 150 μm thickness using a vibratome (Leica).

For analysis of vascular leakage, P6 mice were injected intraperitoneally with Sulfo-NHS-biotin (30 ul of 20 mg/ml Sulfo-NHS-biotin in PBS per mouse) 10-15 minutes before sacrifice. After IP injection, the tracer rapidly equilibrates into the systemic circulation. Mice were deeply anesthetized on ice and perfused via the cardiac route with PBS followed by arachnoid dissection and PFA fixation, as described above.

For cross-sections of isolated dura and arachnoid, fixed tissues were embedded in optimal cutting temperature compound (OCT, Tissue-Tek), rapidly frozen in dry ice, and stored at −80°C. 30 μm sections were cut on a cryostat and thaw-mounted onto Superfrost plus slides. Slides were stored at −80°C until further processing.

### Immunohistochemistry

Whole mount meninges or brain sections were incubated overnight at 4°C with primary antibodies diluted with PBSTC (PBS with 1% Triton X-100, 0.1 mM CaCl2) plus 10% normal goat serum (NGS). Tissues were washed four times with PBSTC over the course of 6–8 hours, and then incubated overnight at 4°C with secondary antibodies diluted in PBSTC plus 10% NGS. If a primary rat antibody was used, secondary antibodies were additionally incubated with 0.5% normal mouse serum as a blocking agent. The following day, tissues were washed at least four times with PBSTC over the course of 6-8 hours, flat-mounted on Superfrost Plus glass slides (Fisher Scientific), and coverslipped with Fluoromount G (EM Sciences 17984-25).

For meninges cross-sections, sections on slides were covered with 2% PFA/PBS at room temperature for 15 minutes, washed three times in PBS, and incubated overnight with primary antibodies diluted in PBSTC plus 10% NGS at 4°C. The following day, sections were washed at least four times with PBSTC and incubated with secondary antibodies for 2 hours at room temperature. Sections were then washed four times with PBSTC and coverslipped with Fluoromount G. For each immunostaining analysis, whole mounts and sections were stained from at least two independent experiments.

### Confocal microscopy

Confocal images were captured with a Zeiss LSM700 confocal microscope (20x and 63x objectives) using Zen Black 2012 software, and processed with Fiji-ImageJ, Adobe Photoshop, and Adobe Illustrator. For Sulfo-NHS-biotin detection with streptavidin, ~2-fold animal-to-animal variability is typically seen in the overall intensity of the strepatvidin signal, most likely due to variable uptake of Sulfo-NHS-biotin from the site of IP injection. To permit a clearer comparison between images of Sulfo-NHS-biotin leakage, this intensity variation has been minimized by manually adjusting the brightness of the Sulfo-NHS-biotin channel.

### Arachnoid tissue lysate

To prepare arachnoid proteins for western blotting, anesthetized mice were perfused with PBS and their brains were dissected in PBS. The arachnoid was detached from the surface of the brain tissue with tweezers and then transferred into a 1.5 mL eppendorf tube and stored at −80°C for further processing. Frozen arachnoids from two mice were pooled, lysed in lysate buffer [50 mM Tris-HCl (pH, 7.4), 150 mM NaCl, 2 mM MgCl_2_, 1% Triton X-100, 0.25 U/μL Benzonase], and then homogenized using a plastic pestle fitted for eppendorf tubes. The homogenates were incubated for 10 minutes at room temperature to digest the nuclear DNA, and then SDS was added to a final concentration of 0.5%. The lysate in SDS was incubated for 20 minutes at 4°C and then centrifuged at 14,000 x g for 15 minutes at 4°C. The supernatant was recovered and its protein concentration was determined using the BCA protein assay kit.

### Immunoblotting

Protein samples (6-8 ug per sample) were loaded onto a 4%-12% NuPAGE Bis-Tris protein gel, which was run at 130 V for 1.5 hours and then blotted onto a nitrocellulose membrane (Millipore). Membranes were blocked with Intercept blocking buffer (LI-COR 927-60001) at room temperature for 1 hour and then probed overnight with primary antibodies diluted in blocking buffer at 4°C. The following day, the membranes were washed four times with TBST and then probed at room temperature for 2 hours with the corresponding infrared secondary antibodies diluted in blocking buffer. Bands were visualized with an Odyssey Fc Imager (LI-COR) and band intensities were quantified with Fiji-Image J software.

### snRNAseq

Two and three independent biological replicate libraries were prepared for the control and *E. coli*-infection groups, respectively, with one P6 mouse used per library. For each sample, the dura and arachnoid were rapidly dissected in ice-cold DPBS (Gibco 14287072). The tissue was minced with a razor blade and Dounce homogenized using a loose-fitting pestle in 5 mL homogenization buffer (0.25 M sucrose, 25 mM KCl, 5 mM MgCl_2_, 20 mM Tricine-KOH,pH 7.8) supplemented with 1 mM DTT, 0.15 mM spermine, 0.5 mM spermidine, EDTA-free protease inhibitor (Roche 11836 170 001), and 60 U/mL RNasin-Plus RNase Inhibitor (Promega N2611). A 5% IGEPAL-630 solution was added to bring the homogenate to 0.3 % IGEPAL CA-630, and the sample was further homogenized with ten strokes of a tight-fitting pestle. The sample was filtered through a 50 μm filter (CellTrix, Sysmex, 04-004-2327), underlayed with solutions of 30% and 40% iodixanol (Sigma D1556) in homogenization buffer, and centrifuged at 10,000×g for 18 minutes in a swinging bucket centrifuge at 4°C. Nuclei were collected at the 30%-40% interface, diluted with two volumes of homogenization buffer, and concentrated by centrifugation for 10 minutes at 500xg at 4°C. snRNAseq libraries were constructed using the 10x Genomics Chromium single-cell 3’ v3 kit following the manufacturer’s protocol (https://support.10xgenomics.com/single-cell-gene-expression/library-prep/doc/user-guide-chromium-single-cell-3-reagent-kits-user-guide-v31-chemistry). Libraries were sequenced on an Illumina NovaSeq 6000.

### Analysis of snRNAseq data

Reads were aligned to the mm10 pre-mRNA index using the Cell Ranger count program, version 3.1.0. The data for the different libraries was merged using the Cell Ranger merge command. Data analysis was performed using the Seurat R package, version 4.0.1 in RStudio. After filtering out nuclei with >1% mitochondrial transcripts or with <500 or >6000 genes, 48,941 nuclei were retained, 14,357 nuclei from the two control samples and 34,585 nuclei from the three infected samples. The data were normalized using a regularized negative binomial regression algorithm implemented in the SCTransform function as described in Hafemeister and Satija, (2019). UMAP dimensional reduction was performed using the R uwot package (https://github.com/jlmelville/uwot) integrated into the Seurat R package. To compare cell types across treatments, the data was integrated using the strategy described in Stuart and Satija (2019). This pipeline involves splitting the dataset by treatment using the Seurat SplitObject function and integrating the subset objects using FindIntegrationAnchors and IntegrateData functions. Data for the various scatter plots were extracted using the Seurat AverageExpression function, and differential gene expression was analyzed using the Seurat FindMarkers function. The Wilcoxon Rank Sum test was used to calculate p-values. The p-values were adjusted with a Bonferroni correction using all genes in the dataset. Data exploration, analysis, and plotting were performed using RStudio (RStudio Team, 2020), the tidyverse collection of R packages (Wickham, 2017), and ggplot2 (Wickham, 2009). For Gene set enrichment analysis (GSEA; Subramanian et al. 2005), genes were ranked by the fold expression change between control and infected datasets. The ranked gene list was used to detect enriches gene sets within the Broad Institute Hallmark Gene Sets using the fgsea R package (https://github.com/ctlab/fgsea).

### Macrophage quantification in the meninges

The density of macrophages was quantified using Fiji-ImageJ (https://imagej.net/software/fiji/) from captured Z-stacked arachnoid or dura wholemounts. The numbers of CD206+PU.1 or CD206+ cells were quantified in representative regions and then normalized to the area analyzed.

### Cell orientation analysis

Using Fiji-ImageJ, the axis of blood flow of the midline vein located in the superior sagittal sinus was defined as 0°, and then the long axis of each EC nucleus, identified by ERG immunostaining, was scored and its angle calculated relative to 0°. Nuclei were analyzed from three independent control vs. infection experiments.

### Fraction of area covered by blood vessels

The relative area covered by arachnoid blood vessels was determined from Z-stacked flatmount images that had been immunostained with CLDN5 or PECAM1. More specifically, flatmount images of representative regions that were populated by capillaries, but not by large veins or arteries, were overlayed in Adobe Illustrator with arrays of ten parallel and evenly-spaced straight white lines. The length of each line corresponds to 166 um on the image and the first and tenth lines in each array are separated by a distance that corresponds to 150 um on the image. Thus, the ten lines define a 166 um x 150 um rectangle. Along each white line, the widths of all of the regions in the image that were not covered by blood vessels were manually scored by drawing (in Adobe Illustrator) a straight line across the vessel-free region. When all of the vessel-free line segments had been drawn for a given square array, the sum of their lengths was calculated with Fiji-ImageJ and divided by the sum of the lengths of the ten white lines to generate an estimate for the fraction of the area not covered by blood vessels. The fraction of the area covered by blood vessels equals one minus the fraction of the area not covered by blood vessels. Each data point in a blood vessel area plot represents the area estimate from one set of white lines, i.e. the estimate obtained from sampling a length of 10 x 166 um = 1.66 mm.

### Statistical analysis

All statistical values are presented as mean ± SD. Statistical tests were carried out using Graphpad Prism 8. The student’s t-test was used to measure statistical significance between two independent groups. One-way analysis of variance (ANOVA) with an appropriate multiple comparisons test was used to compare three or more independent groups. The statistical significance is represented graphically as n.s., not significant (i.e. p>0.05); *, p <0.05; **, p < 0.01; ***, p < 0.001; ****, p < 0.0001.

## Acknowledgements

Supported by the Howard Hughes Medical Institute. The authors thank Dr. Latika Nagpal for helpful comments on the manuscript.

## Supplemental Figure Legends

**Supplemental Figure 1.**
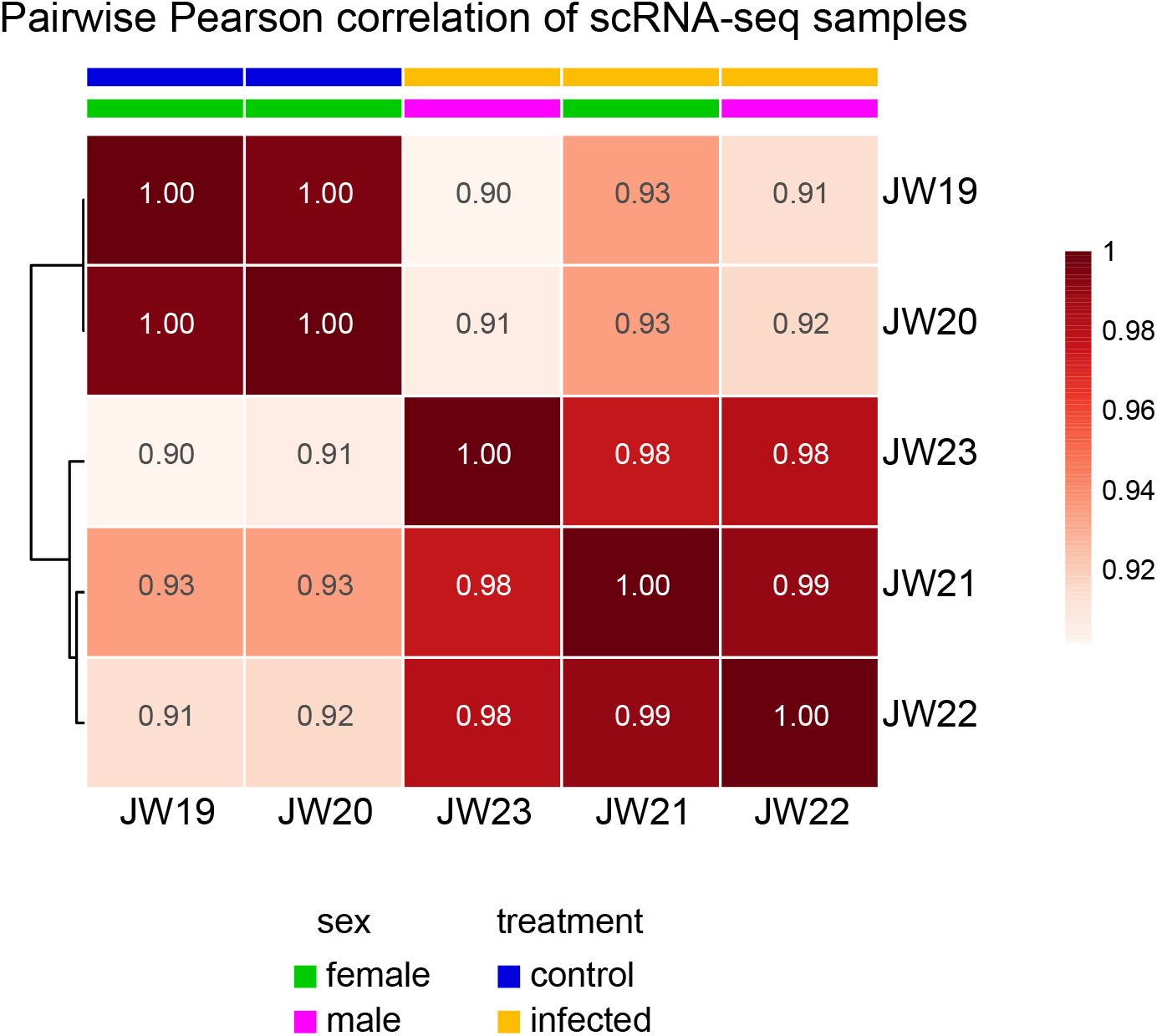
Pairwise Pearson correlations among snRNAseq datasets. Correlation coefficients and hierarchical clustering are shown for two control and three infected snRNAseq datasets from P6 mice. The sex of the mice is indicated.

**Supplemental Figure 2.**
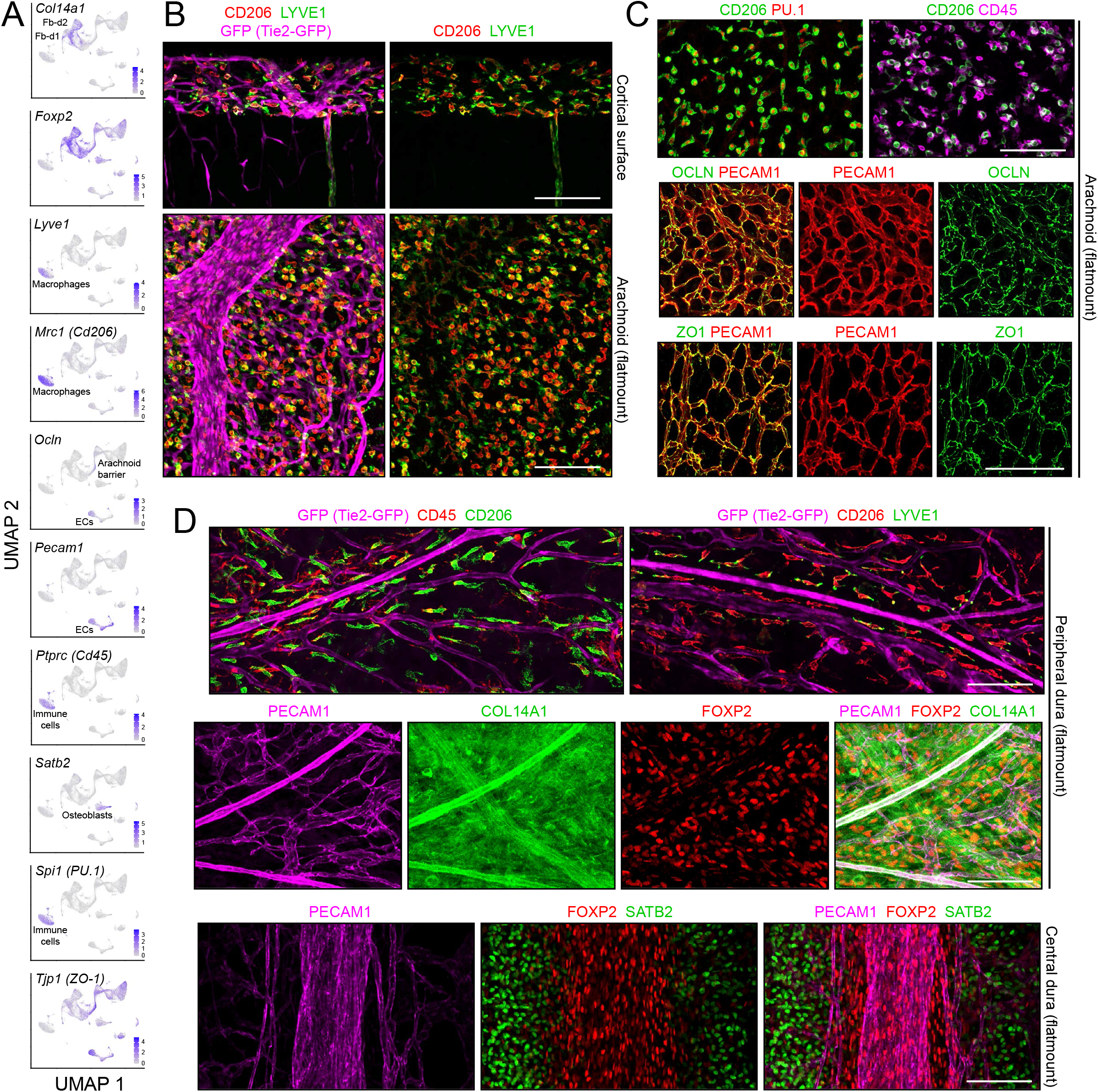
Flatmount images of control arachnoid and dura. (A) snRNAseq UMAP plots of markers used in (B)-(D). (B) Immunostaining for macrophage markers CD206 and LYVE: a comparison between a vibratome cross-section of the cortical surface (upper panels) and an arachnoid flatmount (lower panels). In the upper panels, the cortex occupies the lower ~60% of the image, and the arachnoid occupies most of the upper 40%. (C) Immunostaining of arachnoid flatmounts for macrophage markers CD206 (cytoplasmic/surface), PU.1 (nuclear), and CD45 (cytoplasmic/surface) (upper two panels) and for EC markers OCLN, PECAM1, and ZO1 (lower six panels). (D) Flatmounts of dura, with bone attached, stained for macrophage markers CD206, CD45, and LYVE1 (top panels); EC marker PECAM1 and fibroblast markers COL14A1 (extracellular) and FOXP2 (nuclear) (central panels); and EC marker PECAM1, fibroblast marker FOXP2 (nuclear), and osteoblast marker SATB2 (nuclear) (bottom panel). Scale bars: B-D, 100 um.

**Supplemental Figure 3.**
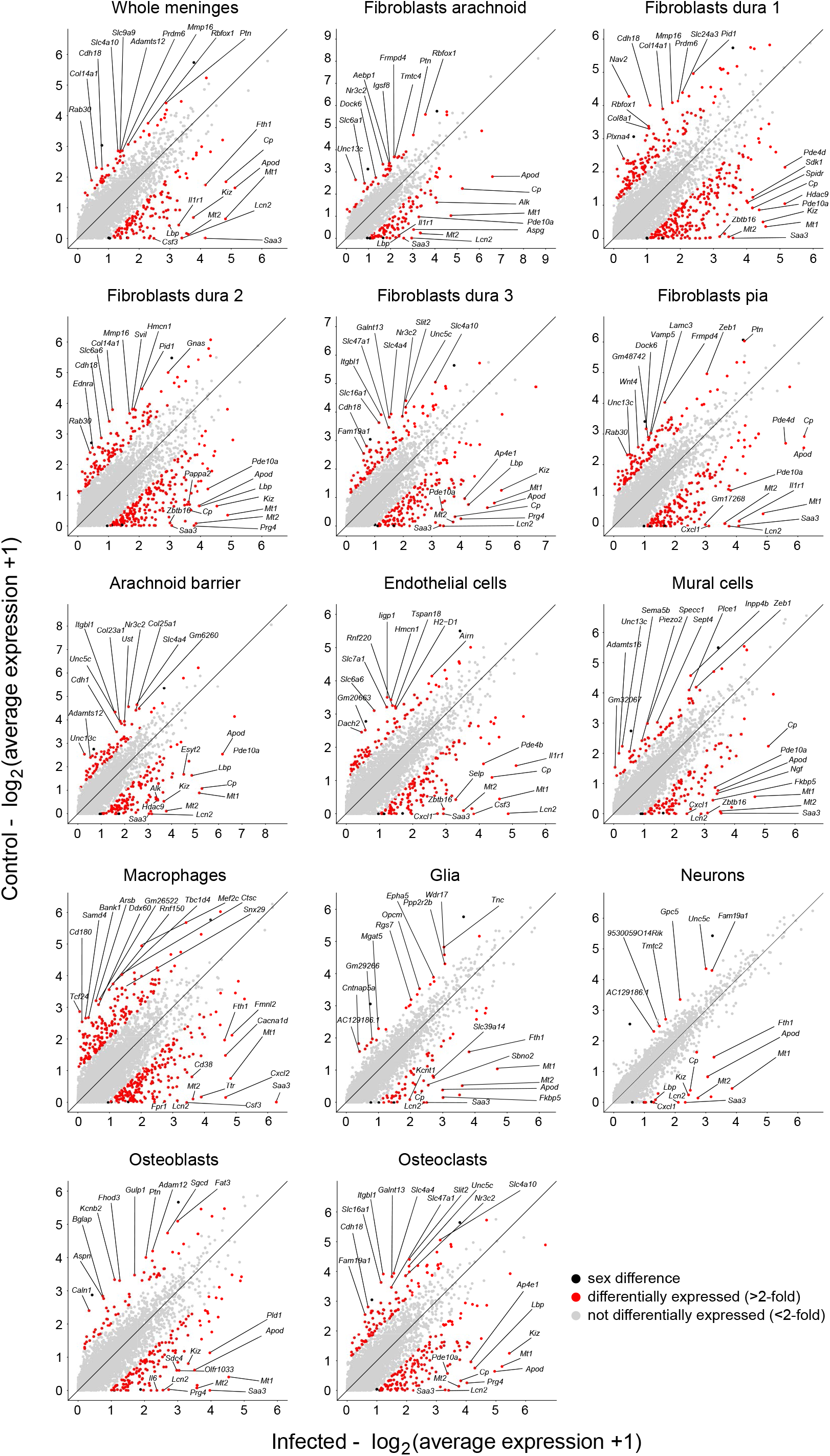
Scatterplots for the major meningeal cell types comparing snRNAseq transcript abundances in control vs. infected mice. Transcripts with a <2-fold abundance difference are shown as light grey dots; transcripts with a >2-fold abundance difference are shown as red dots. Several datapoints with >2-fold abundance difference are due to sex differences between control and infected datasets (e.g. *Xist*); these are shown as unlabeled black dots.

**Supplemental Figure 4.**
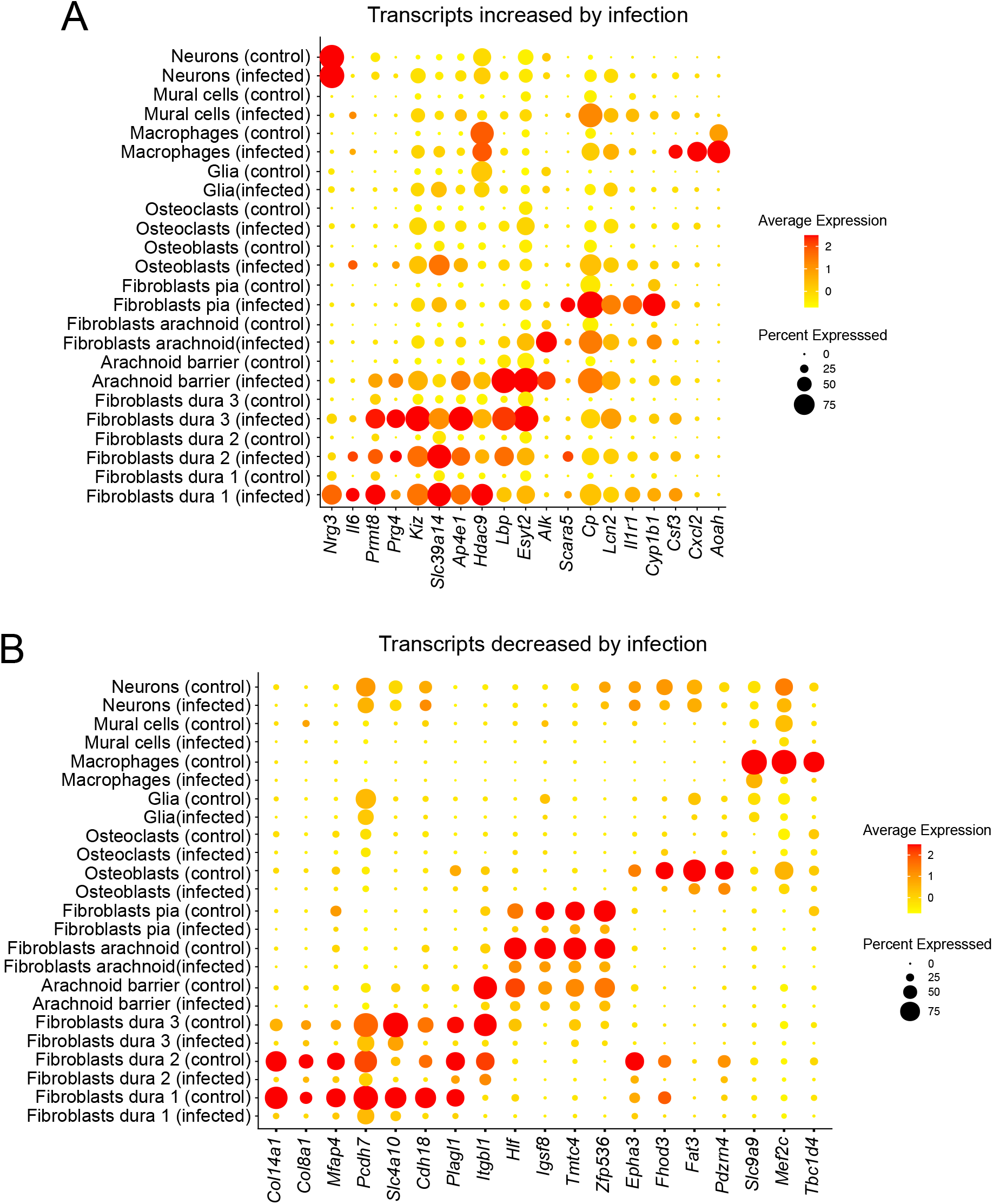
Dot plot showing some of the transcript abundances that distinguish control vs. infected meninges, plotted by cell type. Upper dot plot, transcripts that increase in abundance with infection. Lower dot plot, transcripts that decrease in abundance with infection.

**Supplemental Figure 5.**
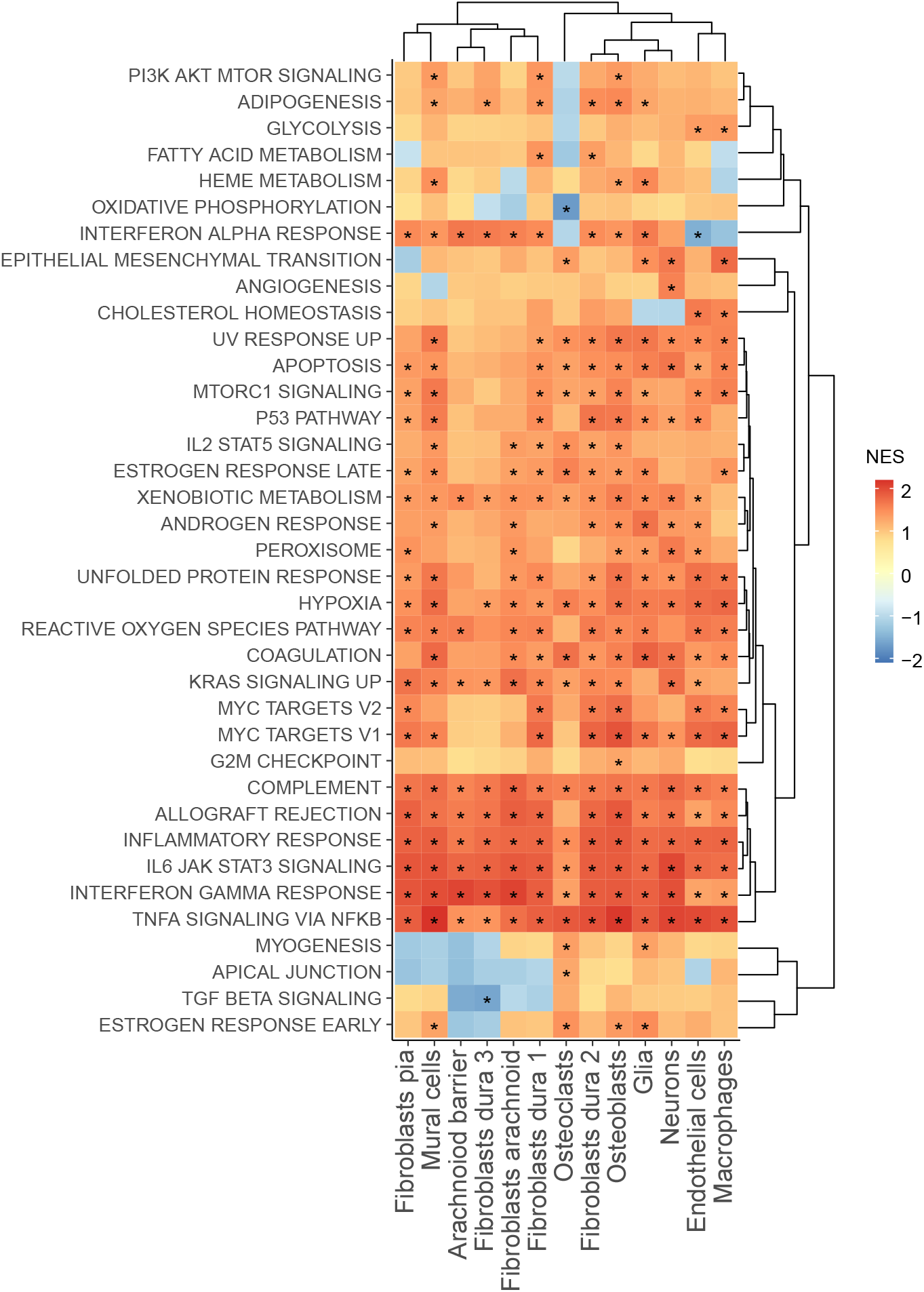
Gene set enrichment analysis (GSEA) for individual cell types in a comparison of control vs. infected meninges. Hierarchical clustering is shown for both gene sets and cells types. Red, transcript abundances increased with infection. Blue, transcript abundances decreased with infection. NES, normalized enrichment score.

**Supplemental Figure 6.**
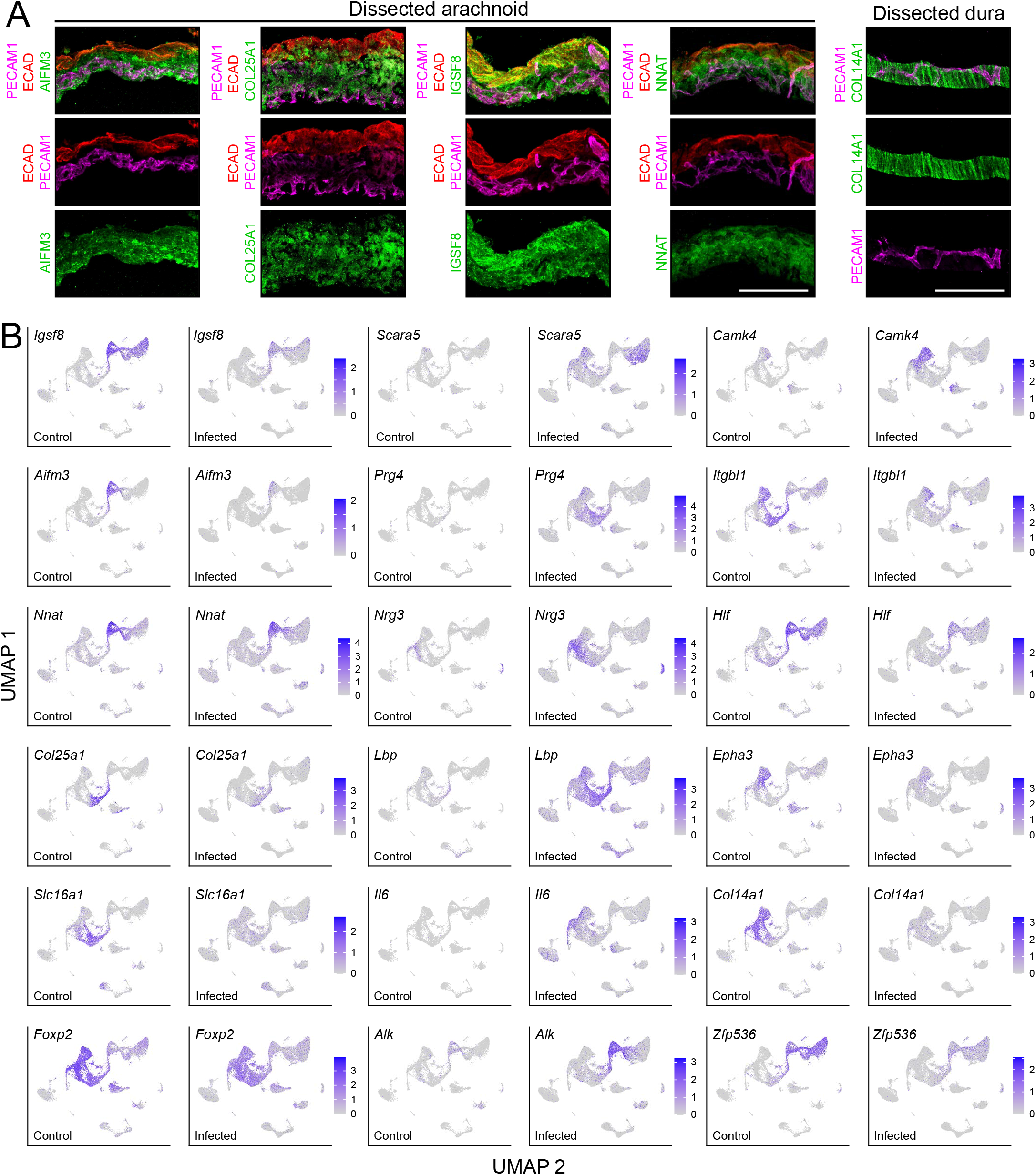
Arachnoid and dura fibroblast markers and fibroblast responses to infection. (A) Cross-sections of dissected arachnoid immunostained for fibroblast markers AIFM3, COL25A1, IGSF8, and NNAT, and of dissected dura immunostained for fibroblast marker COL14A1. (B) UMAP plots from combined control and infected meninges snRNAseq datasets showing changes in abundance in control vs. infected samples for 18 transcripts. Scale bars: A, 100 um.

**Supplemental Figure 7.**
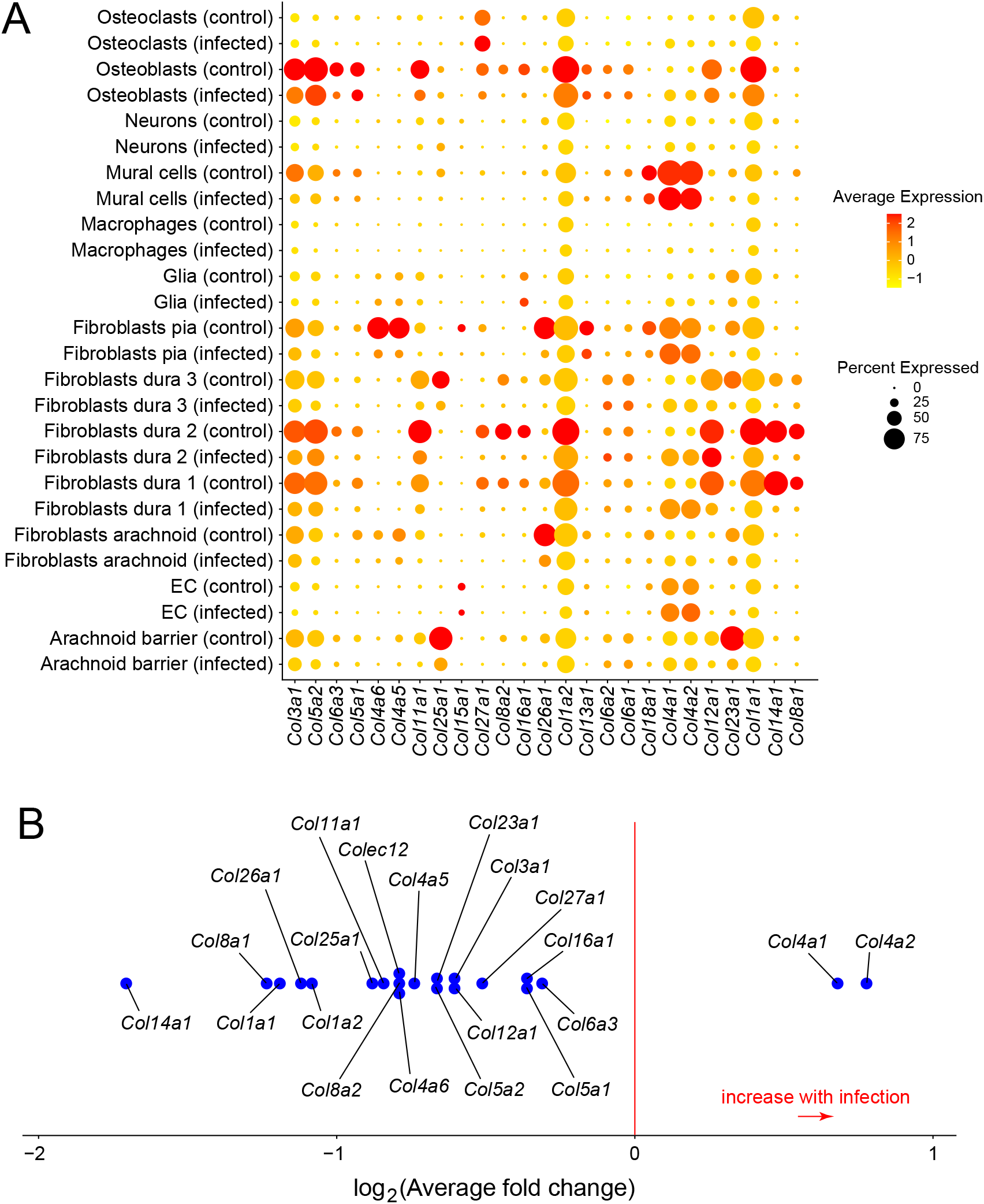
Collagen transcripts in individual meningeal cell types from control vs. infected mice. (A) Dot plot showing abundances for collagen transcripts with detectable expression in the meninges. (B) Fold change in transcript abundances summed over all meninges cell types. Only transcripts with a log_2_-fold change >0.25 are plotted.

**Supplemental Figure 8.**
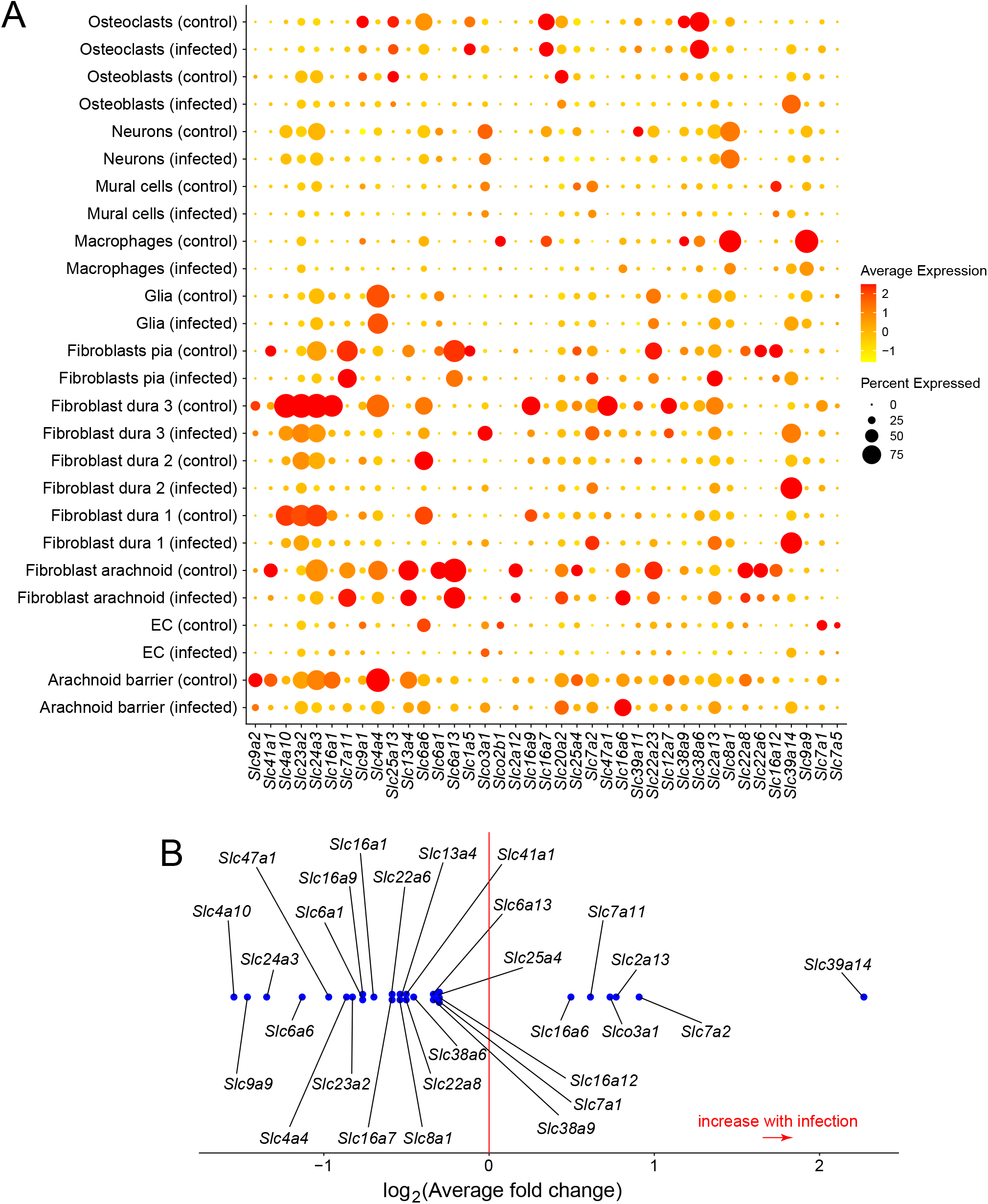
SLC transporter transcripts in individual meningeal cell types from control vs. infected mice. (A) Dot plot showing abundances for SLC transcripts with detectable expression in the meninges. (B) Fold change in transcript abundances summed over all meninges cell types. Only transcripts with a log_2_-fold change >0.25 are plotted.

**Supplemental Figure 9.**
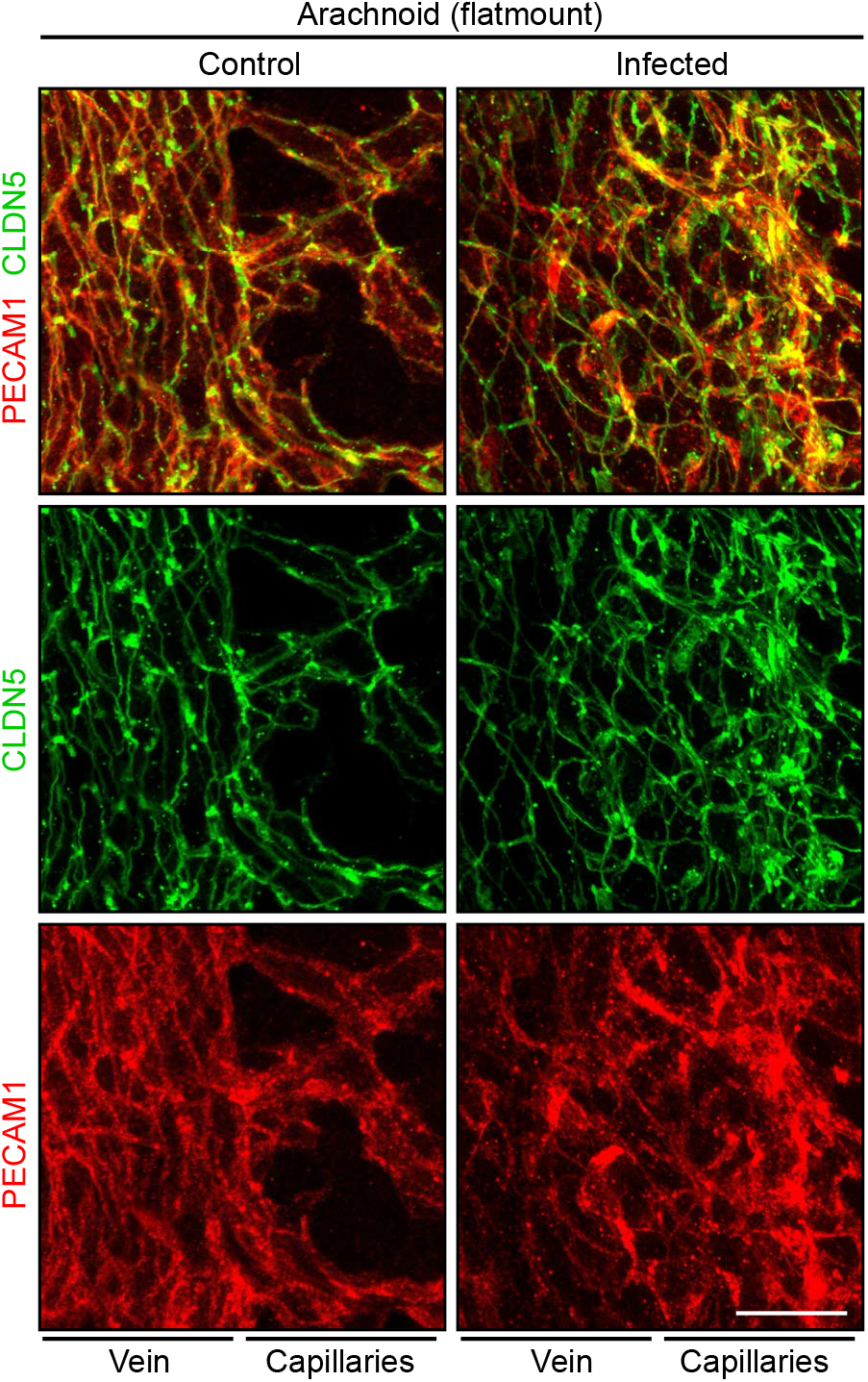
Aberrant EC morphology and subcellular localization of EC proteins in the infected arachnoid. Confocal images captured with a 63x lens show the infection-induced redistribution of CLDN5 and PECAM1 in capillary ECs in arachnoid flatmounts. In each image, a large vein is on the left and several capillaries are on the right. Scale bar: 20 um.

## Supplemental Table Legends

Supplemental Table 1. snRNAseq library statistics.

Supplemental Table 2. Criteria for assigning cell type clusters based on snRNAseq transcript profiles.

Supplemental Table 3. Differential transcript abundances among the major meningeal cell types in infected vs. control snRNAseq datasets. FC, fold change. (Excel File)

Supplemental Table 4. Differential transcript abundances among CCL2^-^ and CCL2^+^ resident macrophages in infected vs. control snRNAseq datasets. FC, fold change. (Excel File)

Supplemental Table 5. Differential transcript abundances among EC cell types in infected vs. control snRNAseq datasets. FC, fold change. (Excel File)

---

**Supplemental Table 1.**

| library | treatment | number<br>of nuclei | mean number of<br>transcripts per nucleus |
| --- | --- | --- | --- |
| JW19 | uninfected | 7282 | 1680.02 |
| JW20 | uninfected | 7074 | 1618.53 |
| JW21 | infected | 8521 | 1476.26 |
| JW22 | infected | 11195 | 1232.82 |
| JW23 | infected | 14869 | 976.33 |
| total |  | 48941 | 1320 |

**Number of nuclei per cluster**
| Cell type | infected | control | total |
| --- | --- | --- | --- |
| Arachnoid barrier cells | 1236 | 396 | 1632 |
| Endothelial cells | 3558 | 1154 | 4712 |
| Fibroblasts, arachnoid | 4043 | 2111 | 6154 |
| Fibroblasts, dura 1 | 4280 | 1798 | 6078 |
| Fibroblasts, dura 2 | 2767 | 1647 | 4414 |
| Fibroblasts, dura 3 | 2857 | 1304 | 4161 |
| Fibroblasts, mitotic | 668 | 312 | 980 |
| Fibroblasts, pia | 5380 | 2199 | 7579 |
| Glia | 2103 | 377 | 2480 |
| Immune cells | 3790 | 1555 | 5345 |
| Mural cells | 437 | 242 | 679 |
| Neurons | 580 | 131 | 711 |
| Osteoblasts | 2010 | 923 | 2933 |
| Osteoclasts | 148 | 66 | 214 |
| U1 | 468 | 119 | 587 |
| U2 | 260 | 22 | 282 |
| total | 34585 | 14356 | 48941 |

**Supplemental Table 2.** List of literature-based markers for characterization of cell identity of meningeal clusters.

| Cell type | Gene | Ref. |
| --- | --- | --- |
| **Myeloid cells/Immune cells** | *Ptprc (Cd45)* | 1, 2 |
| Macrophages | *Csf1r* | 3 |
|  | *C1qa* | 3 |
|  | *F13a1* | 4 |
|  | *Adgre1* | 3, 5 |
|  | *Mrc1/Cd206* | 3, 6 |
|  | *Lyve1* | 3, 6 |
|  | *Apoe* | 3 |
|  | *Spi1* | 1 |
| CCL2+ macrophages | *Ccl2* | 2 |
| Monocytes/Monocytes-derived cells | *Ccr2* | 6, 7 |
|  | *H2-d1* | 8 |
|  | *Ly6c2* | 6, 9 |
|  | *S100a8* | 10 |
|  | *S100a9* | 10 |
|  | *Cytip* | 11, 12 |
| ILC2 | *Il7r* | 13 |
|  | *Gata3* | 13 |
|  | *Il1rl1* | 13 |
| ILC3 | *Rorc/Rorgt* | 13 |
| T cells | *Cd3e* | 2 |
|  | *Cd4* | 2 |
| Microglia | *Ccr5* | 14 |
|  | *Siglech* | 15 |
|  | *Hexb* | 16 |
|  | *P2ry12* | 17 |
| Osteoclasts | *Ctsk* | 18 |
|  | *Atp6v0d2* | 19, 20 |
|  | *Igtb3* | 21 |
| Endothelial cells | *Pecam1* | 22, 23 |
|  | *Erg* | 24 |
|  | *Tek* | 25 |
| Endothelial cell, arachnoid | *Cldn5* | 22, 26 |
|  | *Lef1* | 27, 28 |
|  | *Slc7a5* | 22 |
|  | *Lrp8* | 29 |
|  | *Slco1c1* | 30 |
| Endothelial cell, dura | *Plvap* | 31, 32 |
| Endothelial cell, dura, VWF+ | *Vwf* | 32, 33, 34 |
| Endothelial cell, artery | *Bmx* | 22, 35 |
|  | *Fbln5* | 23 |
|  | *Vegfc* | 22, 36 |
| **Mural cells** | *Abcc9* | 32 |
|  | *Notch3* | 37 |
|  | *Pdgfrb* | 32 |
| **Fibroblasts** | *Col1a1* | 38 |
|  | *Pdgfra* | 22, 38 |
| Fibroblast, pia | *Lama1* | 38 |
|  | *Cxcl12* | 38 |
|  | *Col15a1* | 39 |
| Fibroblast, arachnoid | *Nnat* | 38 |
|  | *Aldh1a2* | 38 |
|  | *Ptgds* | 38 |
| Fibroblast, dura | *Fxyd5* | 38 |
|  | *Foxp1* | 40 |
|  | *Mgp* | 38 |
| Arachnoid barrier | *Cdh1* | 38, 41 |
|  | *Cldh11* | 38 |
|  | *Tjp1* | 38 |
| **Osteoblasts** | *Runx2* | 42 |
|  | *Alpl* | 43 |
| **Neurons** | *Rbfox3* | 44 |
|  | *Nrxn3* | 45 |
| **Glia** | *Gfap* | 46 |
|  | *Aqp4* | 47 |
| Mitotic cells | *Top2a* | 48 |
|  | *Mki67* | 49 |

